# *NvPOU4/Brain3* functions as a terminal selector gene in the nervous system of the cnidarian *Nematostella vectensis*

**DOI:** 10.1101/2020.01.08.898437

**Authors:** Océane Tournière, David Dolan, Gemma Sian Richards, Kartik Sunagar, Yaara Y Columbus-Shenkar, Yehu Moran, Fabian Rentzsch

## Abstract

Terminal selectors are transcription factors that control the morphological, physiological and molecular features that characterize distinct cell types. Here we use expression analyses and a transgenic reporter line to show that *NvPOU4* is expressed in post-mitotic cells that give rise to a diverse set of neural cell types in the sea anemone *Nematostella vectensis*. We generated a loss-of-function allele by CRISPR/Cas9 and used additional transgenic reporter lines to show that the initial specification of neural cells is not affected in the *NvPOU4* mutants. Analyses of transcriptomes derived from the mutants and from different neural cell populations revealed that *NvPOU4* is required for the execution of the terminal differentiation program of these neural cells. These findings suggest that POU4 genes have ancient functions as terminal selectors for morphologically and functionally highly disparate types of neurons and they provide experimental support for the relevance of terminal selectors for understanding the evolution of cell types.

## INTRODUCTION

Neurons display a remarkable morphological and molecular diversity. The acquisition of the features that characterize different types of neurons is the result of a series of developmental processes largely directed by transcription factors and signalling molecules (Edlund and Jessell, 1999). Early stages of neural development are often characterized by the proliferation of different types of progenitor cells via symmetric and/or asymmetric divisions (Doe, 2008; Homem et al., 2015; Taverna et al., 2014). After their terminal mitosis, the differentiation of neurons typically begins with the occurrence of more general neural features like the expression of neural cytoskeletal proteins and by the formation of neurites (Ernsberger, 2012; Stefanakis et al., 2015). The terminal identity of individual types of neurons eventually manifests by the expression of specific neurotransmitter systems, the elaboration of specific projection patterns and other factors defining the physiological properties of these neuron types. Transcription factors that regulate these terminal differentiation features of distinct neuron types are called terminal selector genes (Allan and Thor, 2015; Hobert, 2016; Hobert and Kratsios, 2019). Terminal selectors often function in combination and may affect all or only some aspects of the identity of a neuron (e.g. (Etchberger et al., 2007; Stratmann et al., 2019). While transcription factors regulating the terminal differentiation of neurons have been identified in several bilaterians (Allan and Thor, 2015; Hobert and Kratsios, 2019), it is currently unknown whether conserved terminal selectors tend to have comparable functions over long evolutionary distances. We address this question here by analysing the function of the POU domain transcription factor *POU4/Brn3* in a representative of a non-bilaterian animal clade, the cnidarian *Nematostella vectensis*.

Cnidarians are the sister group of bilaterians (Dunn et al., 2014; Telford et al., 2015), with the separation of these two lineages estimated to have occurred over 600 million years ago (dos Reis et al., 2015; Park et al., 2012). As adults, they possess a relatively simple nervous system that lacks brain-like centralization. Their nervous system comprises three main classes of neural cells: cnidocytes (“stinging cells”, cnidarian-specific mechano/chemoreceptor cells), ganglion cells (interneuron-like cells) and sensory/sensory-motor cells (Galliot et al., 2009; Rentzsch et al., 2019; Watanabe et al., 2009). Morphological and molecular analyses suggest the existence of distinct subpopulations of these classes of neural cells; however, an integrated characterization of neural cell types in cnidarians is currently lacking (Rentzsch et al., 2019; Sebe-Pedros et al., 2018; Siebert et al., 2018). The sea anemone *Nematostella vectensis* belongs to the anthozoan class of cnidarians. Due to its inducible fertilization, its relatively short generation time and amenability to molecular manipulations, *Nematostella* has become an important cnidarian model organism (Layden et al., 2016). It has previously been shown that a large fraction of its neurons derives from a pool of *NvSoxB(2)*-expressing neural progenitor cells (NPCs) located in both ectoderm and endoderm, which give rise to the three classes of neural cells (Nakanishi et al., 2012; Richards and Rentzsch, 2014). In addition to *NvSoxB(2)*, the basic helix-loop-helix (bHLH) genes *NvAshA* and *NvAtonal-like* have been identified as positive regulators of neurogenesis, whereas Notch signalling acts to restrict the number of neural progenitor cells (Layden et al., 2012; Layden and Martindale, 2014; Rentzsch et al., 2017; Richards and Rentzsch, 2015). For the regulation of early stages of neuron development, these observations suggest a considerable degree of conservation between *Nematostella* and bilaterians. How the terminal differentiation of neurons is regulated in *Nematostella*, is currently poorly understood. Accordingly, it is not known whether the conservation of neurogenic transcriptional programs between *Nematostella* and bilaterians extends to the late stages of neuron development.

POU (Pit/Oct1/UNC-86) genes are transcription factors that contain a bipartite DNA binding domain consisting of a POU-specific and a homeobox domain. While being found only in metazoans, they diversified early during animal evolution and four classes of POU genes were present in the last common ancestor of all living animals (Gold et al., 2014; Larroux et al., 2008). Genes of the POU4 class are predominantly expressed in neuronal cells and have been shown to regulate the terminal differentiation of these neurons in several organisms. In mammals there are three POU4 genes, *Brn3a*, *Brn3b* and *Brn3c,* all of which are prominently expressed in partially overlapping areas of sensory structures, as well as in other parts of the nervous system (Collum et al., 1992; Fedtsova and Turner, 1995; Gerrero et al., 1993; Ninkina et al., 1993; Turner et al., 1994; Xiang et al., 1995; Xiang et al., 1993). Analyses of knock-out mice have identified key roles for these genes in the formation of hair cells in the auditory and vestibular systems (*Brn3c*; (Erkman et al., 1996; Xiang et al., 1997), of retinal ganglion cells (*Brn3b*, (Erkman et al., 1996; Gan et al., 1996) and of somatosensory and brainstem neurons (*Brn3a*, (McEvilly et al., 1996; Xiang et al., 1996). Each of the Brn3 genes functions mainly at later stages of neural differentiation, e.g. in the acquisition of morphological features of somatosensory neurons and retinal ganglion cells (Badea et al., 2009; Badea et al., 2012; Erkman et al., 2000; Ryan and Rosenfeld, 1997). In *Drosophila*, the POU4 orthologue *acj6/I- POU* regulates synaptic targeting in the central nervous system and the odour sensitivity of olfactory neurons (Ayer and Carlson, 1991; Certel et al., 2000; Clyne et al., 1999; Treacy et al., 1992). In the nematode *Caenorhabditis elegans*, the single POU4 gene, *unc-8*6, is expressed in several types of neurons (Finney and Ruvkun, 1990). In most of these neurons, *unc-86* acts in specific combinations with other transcription factors to control a terminal differentiation program, for example by defining the neurotransmitter identity of the cell (Chalfie et al., 1981; Duggan et al., 1998; Hobert, 2016; Serrano-Saiz et al., 2013; Zhang et al., 2014). This terminal selector function of POU4 genes is often also required for the maintenance of the identity of these neurons, both in *C. elegans* and in mice (Serrano-Saiz et al., 2018). In addition to its role in terminal differentiation, *unc-86* has a role in regulating the division of some neural progenitor cells (Chalfie et al., 1981; Finney and Ruvkun, 1990). In line with potential roles in the nervous system, POU4 genes are expressed in sensory and other neural structures in several other bilaterians (Backfisch et al., 2013; Candiani et al., 2006; Candiani et al., 2005; Nomaksteinsky et al., 2013; O’Brien and Degnan, 2002; Ramachandra et al., 2002; Wollesen et al., 2014). Overall, their roles in different types of neurons and in several bilaterians make POU4 genes prime candidates for addressing the early evolution of terminal neural differentiation.

Outside bilaterians, little is known about the role of POU4 genes. In medusae of the cnidarians *Aurelia sp*. and *Craspedacusta sowerbyi, POU4* expression was detected in sensory structures at the margin of the bell (Hroudova et al., 2012; Nakanishi et al., 2010), however, no functional analyses have been reported so far in these groups. In this study, we use gene expression analyses and an *NvPOU4*::memGFP transgenic reporter line to show that the single *Nematostella* POU class 4 gene is expressed in a large and heterogeneous population of post-mitotic neural cells. Furthermore, we generated an *NvPOU4* mutant line by CRISPR/Cas9- mediated genome editing, analysed the transcriptome of the mutants and crossed it to nervous system-specific transgenic reporter lines. This revealed that *NvPOU4* functions in the terminal differentiation of neural cells, including the cnidarian-specific cnidocytes. These observations indicate that POU4 genes have ancient roles in terminal neural differentiation and that the regulation of cell differentiation by terminal selector genes evolved early in animal evolution.

## RESULTS

### *NvPOU4* is expressed in neural cells from early blastula to polyp stage

The *Nematostella* genome contains a single POU4 gene (Putnam et al., 2007; Larroux et al., 2008; Gold et al., 2014). Using whole mount *in situ* hybridization, we first observed expression of *NvPOU4* in few cells at early blastula (12 hours post fertilization at 21°C; Figure 1A). This expression occurs after the start of *NvSoxB(2)* expression (a gene expressed in neural progenitor cells, (Magie et al., 2005; Richards and Rentzsch, 2014)), but before expression of *NvNCol3* commences (a gene expressed in differentiating cnidocytes and encoding the minicollagen structural protein of the cnidocyst capsule wall (Babonis and Martindale, 2017; Zenkert et al., 2011)). *NvPOU4* is expressed in scattered cells all over the ectoderm of the embryo at gastrula stage and in scattered single cells in both ectoderm and endoderm at mid-planula stage (Figure 1B-C). At late planula stage, the expression is prominent in cells close to the oral opening (Figure 1D). At tentacle bud stage, this expression has resolved into four distinct patches, the developing tentacle buds (Figure 1E). In primary polyps, expression of *NvPOU4* is still detectable in scattered ectodermal and endodermal cells and, most prominently, in the tentacle tips (Figure 1F). The expression in scattered cells throughout the body column resembles that of several neural genes described previously (Layden et al., 2012; Marlow et al., 2009; Nakanishi et al., 2012) whereas the expression in the tentacle buds and tentacle tips reflects the main sites of cnidocyte formation (Babonis and Martindale, 2017; Zenkert et al., 2011).

**Fig. 1.**
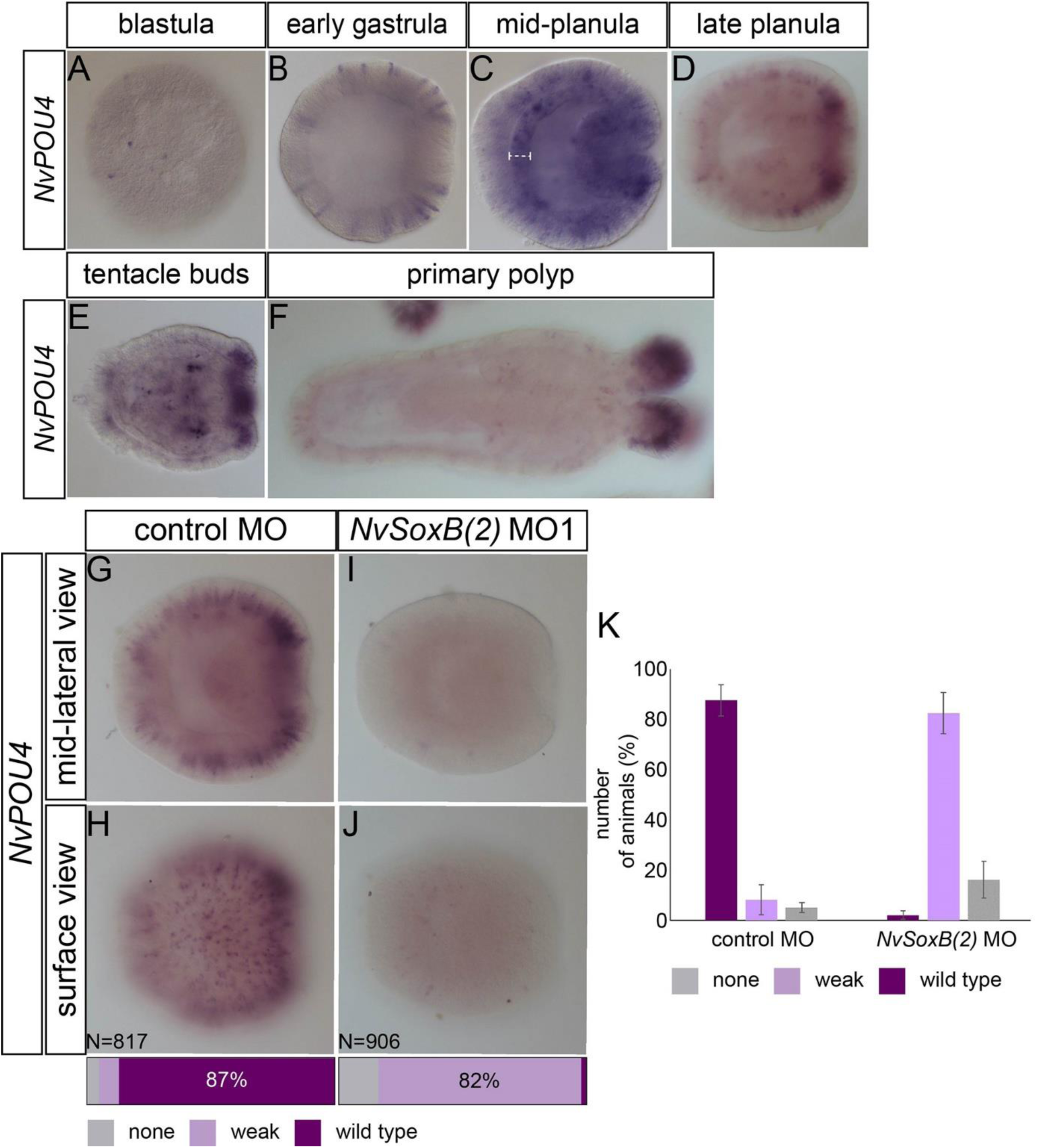
*NvPOU4* expression is controlled by *NvSoxB(2).* (A-F) *In situ* hybridization with probes indicated on the left side and the developmental stage on top. Mid-lateral views with the aboral pole to the left. The white bracket in (C) indicates the endoderm layer. *NvPOU4* is expressed in scattered single cells all over the embryos. (G-J) Treatments are indicated to the top of each image, each MO condition is compared to control MO injected animals. *NvSoxB(2)* MO injection results in a decreased number of *NvPOU4*-expressing cells. Animals were quantified into phenotypic classes based on having no, weak or wild type expression. (I–J) are examples of weak expression. Bars at the base of each image represent the percentage of animals in each phenotypic class. (K) Graphical representation of the percentage of animals in each phenotypic class with standard deviation (four biological replicates).

*NvPOU4* starts being expressed approximately two hours after the expression of *NvSoxB(2)* (Figure S1), a gene that is broadly required for neurogenesis in *Nematostella* (Richards and Rentzsch, 2014). To understand better whether *NvPOU4* is expressed in cells of the neural lineage, we inhibited the function of *NvSoxB(2)* by injection of a morpholino antisense oligonucleotide (Richards and Rentzsch, 2014). This resulted in a nearly complete suppression of *NvPOU4* expression (Figure 1G-K).

Taken together, the expression pattern and the dependence on *NvSoxB(2)* suggest that *NvPOU4* is expressed in neural cells in *Nematostella*.

### *NvPOU4* expression is restricted to non-proliferating, differentiating cells

To better characterize the identity of the *NvPOU4*-expressing cells, we used double fluorescent *in situ* hybridization to test co-expression of *NvPOU4* with other genes expressed during neural development. We observed that *NvPOU4* is partially co-expressed with the neuropeptide gene *NvRFamide* (labelling differentiating sensory and ganglion cells, Figure 2A-B) and with *NvNcol3* (Figure 2C-D). In contrast, *NvPOU4* is not co-expressed with the neural progenitor marker *NvSoxB(2)* from blastula to late planula stages (Figure 2E-F, Figure S2 and Movies S1-S9). As we have shown that the expression of *NvPOU4* depends on *NvSoxB(2)* function (Figure 1G-K), the lack of co-expression of the two mRNAs indicates that they might be expressed sequentially in developing neurons. As a first step to test this possibility, we examined cell proliferation in *NvPOU4* expressing cells. We incubated wild type animals at late blastula stage for 30 min with EdU, fixed them immediately afterwards and performed EdU detection together with fluorescent *in situ hybridization* (Figure 2G-I and Movie S10). We did not observe any EdU positive *NvPOU4*-expressing cells (10 embryos, in total 310 *NvPOU4* expressing cells in 100 μm × 100 μm squares in the mid-lateral part of the blastoderm). This differs from previous observations of EdU-incorporating cells that express *NvSoxB(2)* mRNA (Richards and Rentzsch, 2014).

**Fig. 2.**
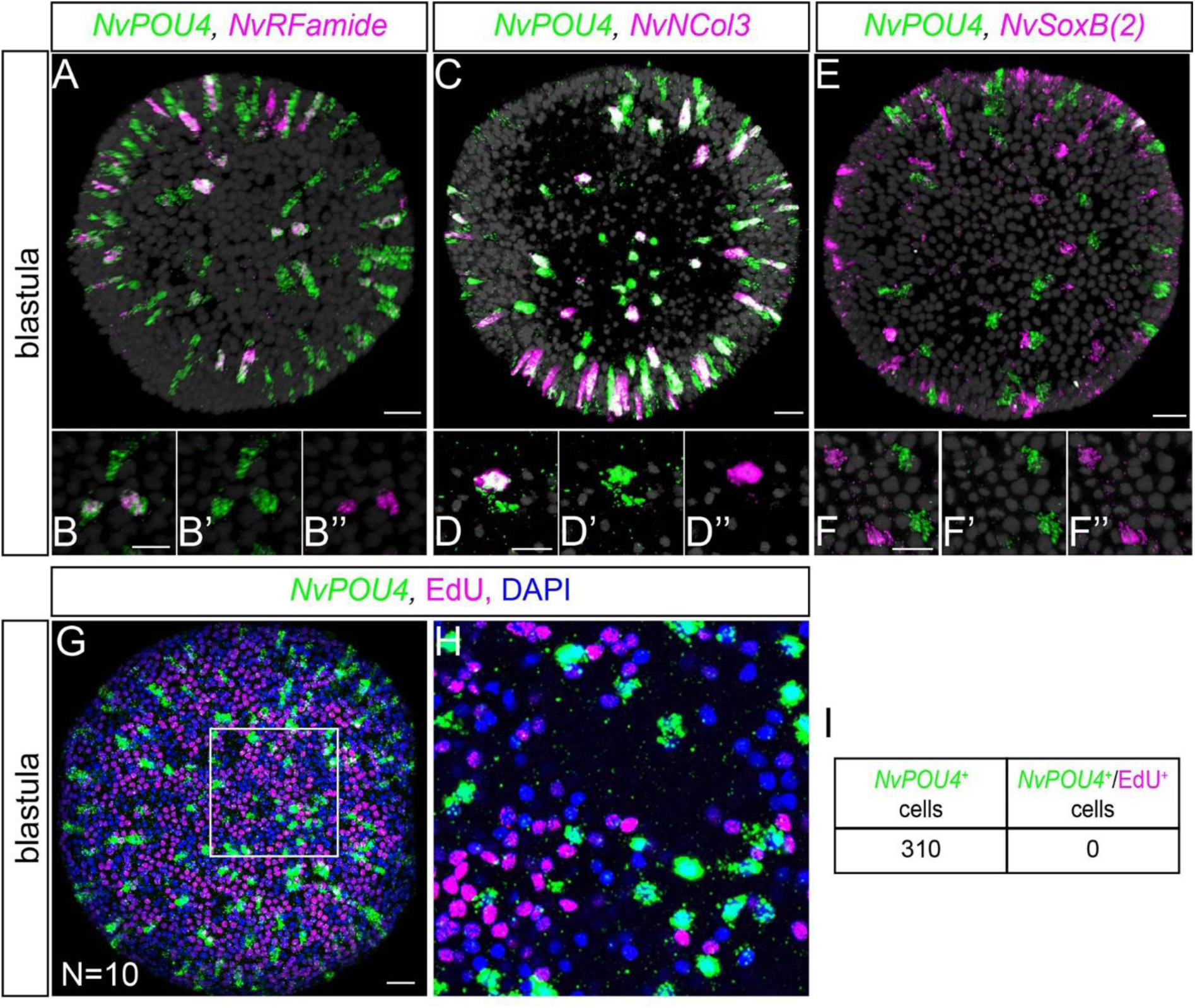
*NvPOU4* is expressed in differentiating cells. (A-F) Double fluorescent *in situ* hybridization at blastula stage for *NvPOU4* (green) and either *NvRFamide, NvNcol3* or *NvSoxB(2)* (magenta). DAPI is shown in grey. The neural differentiating markers *NvRFamide* (A-B) and *NvNCol3* (C-D) show partial co-expression (white) with *NvPOU4*. No co-expression with the NPC marker *NvSoxB(2)* was detected (E-F). (A, C, E) are projections of stacks, all other images are single confocal sections. Stacks are available as Movies S1-3. White color in (E) is caused by the maximal projection and does not represent co-labelling of *NvPOU4* and *NvSoxB(2)*. For gastrula and planula stages, see Figure S2 and Movies S4-9. All *in situ* hybridizations were performed with at least three replicates. (G-H) Fluorescent *in situ* hybridization for *NvPOU4* (green) plus staining for EdU (magenta) and DAPI (blue) at blastula stage. (I) In a 100 μm × 100 μm area of mid-lateral ectoderm, of 310 *NvPOU4*-expressing cells (in 10 animals), none incorporated EdU, suggesting that *NvPOU4*-expressing cells are post-mitotic. (G) is a projections of a stack, (H) is a single confocal section. The stack for (G) is available as Movie S10. Scale bars represent 20 μm

Together, these data suggest that *NvPOU4* is expressed in non-proliferating, differentiating cells of the developing nervous system.

### *NvPOU4* expressing cells develop into neurons and cnidocytes

To gain further insight into the nature of the *NvPOU4*-expressing cells, we generated a stable transgenic reporter line, in which a 4.7 kb upstream region of the *NvPOU4* coding sequence drives the expression of a membrane-tethered GFP (*NvPOU4*::memGFP). This allowed the identification of the *NvPOU4*-expressing cells and their progeny. Double fluorescent *in situ hybridization* of *memGFP* and *NvPOU4* in transgenic embryos, showed a strong co-expression of the reporter gene transcripts and endogenous *NvPOU4* (Figure S3); this confirms that the reporter line accurately reflects the endogenous expression of *NvPOU4*.

Analysis of memGFP expression showed that it starts in early gastrula and is increased and maintained until polyp stages (Figure 3A-C). At early planula stage, memGFP is localized in scattered ectodermal cells that have a slender shape and often an apical cilium (Figure 3A). Later on, at late planula stage, it is possible to detect the memGFP protein in both scattered ectodermal and endodermal cells (Figure 3B). At primary polyp stage, memGFP localization highlights the nerve net and is expressed in cells with various morphologies in the ectoderm and in the endoderm. In the endoderm, neurites of many memGFP^+^ neurons extend along the mesenteries - the longitudinal in-foldings of the endoderm. At this stage, there is also expression of the memGFP protein in scattered ectodermal cells all over the body column and in the tentacles of the animals, with particularly strong labelling in the tips of the tentacles. This pattern of transgene expression is maintained in juvenile and adult polyps (not shown).

**Fig. 3.**
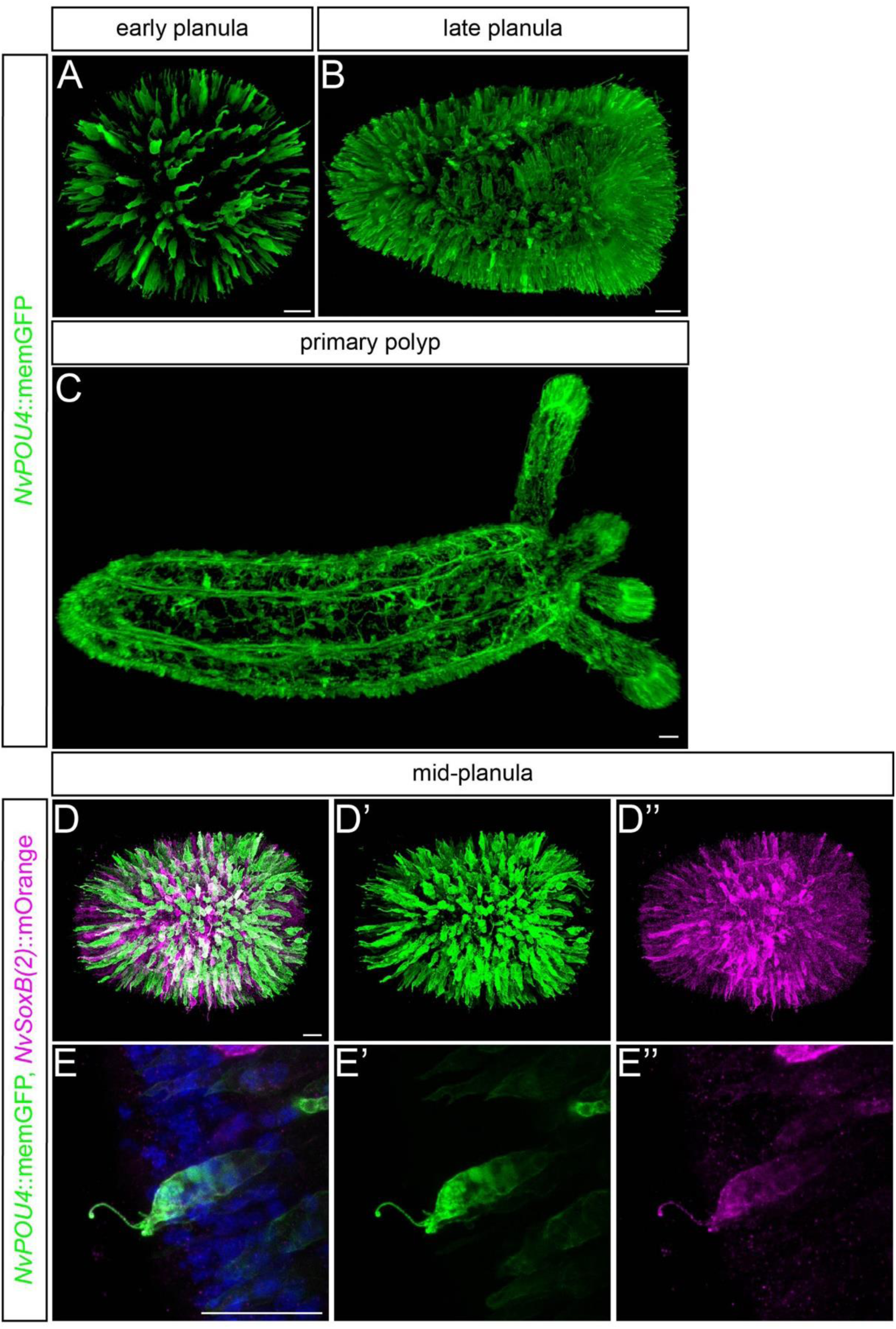
A transgenic reporter line recapitulates the expression of *NvPOU4*. (A-C) Confocal microscopy images of the *NvPOU4*::memGFP transgenic line, memGFP is detected by anti-GFP antibody (green). All images are lateral views with the aboral pole to the left. All images are Imaris snapshots from the 3D reconstructions. Expression of memGFP is detected from gastrula stage on and is consistent with the *in situ* hybridization signals (Figure S3). (D-E) Double transgenic animals with *NvPOU4*::memGFP (green) and *NvSoxB(2)*::mOrange2 (magenta). Lateral views with aboral pole to the left. At planula stage the *NvPOU4*::memGFP^+^ cells are also mOrange^+^. (E-E’’) shows a double-positive putative sensory cell in the ectoderm. Scale bars represent 20 μm

Next, we generated double transgenic animals by crossing the *NvPOU4*::memGFP line to other previously characterized neuronal reporter lines. To clarify the relationship between *NvSoxB(2)* and *NvPOU4* expressing cells, we generated *NvSoxB(2)::memOrange*; *NvPOU4::memGFP* double transgenics. At both gastrula and planula stage, nearly all *NvPOU4*::memGFP cells were also labelled with the *NvSoxB(2)* reporter transgene (Figure 3 D-E). This supports the scenario in which the two genes are expressed sequentially in the same cells, with *NvSoxB(2)* expression preceding that of *NvPOU4*. We also noted that the overlap of the two transgenes is not absolute: while the *NvSoxB(2)::memOrange* transgene is expressed more broadly, there are also some cells that are positive for *NvPOU4*::memGFP, but not for *NvSoxB(2)*::memOrange (Fig. 3D).

The previously characterized *NvNcol3::*mOrange2 line labels differentiating cnidocysts - the extrusive capsules of the cnidocytes (Sunagar et al., 2018). In double transgenic animals (*NvPOU4*::memGFP, *NvNcol3::*mOrange2) (Figure 4A-F), GFP positive membranes surround all mOrange2 positive cnidocysts throughout the animal from mid-planula to polyp stage. Most of the ectodermal memGFP^+^ cells appear to contain a cnidocyst and are therefore differentiating or differentiated cnidocytes (Figure 4B, E, F). However, this co-expression is not absolute, and some memGFP^+^ cells do not contain a developing cnidocyst and have a long apical cilium (Figure 4C). The *NvPOU4* transgene is thus expressed in developing cnidocytes and potentially other cell types.

**Fig. 4.**
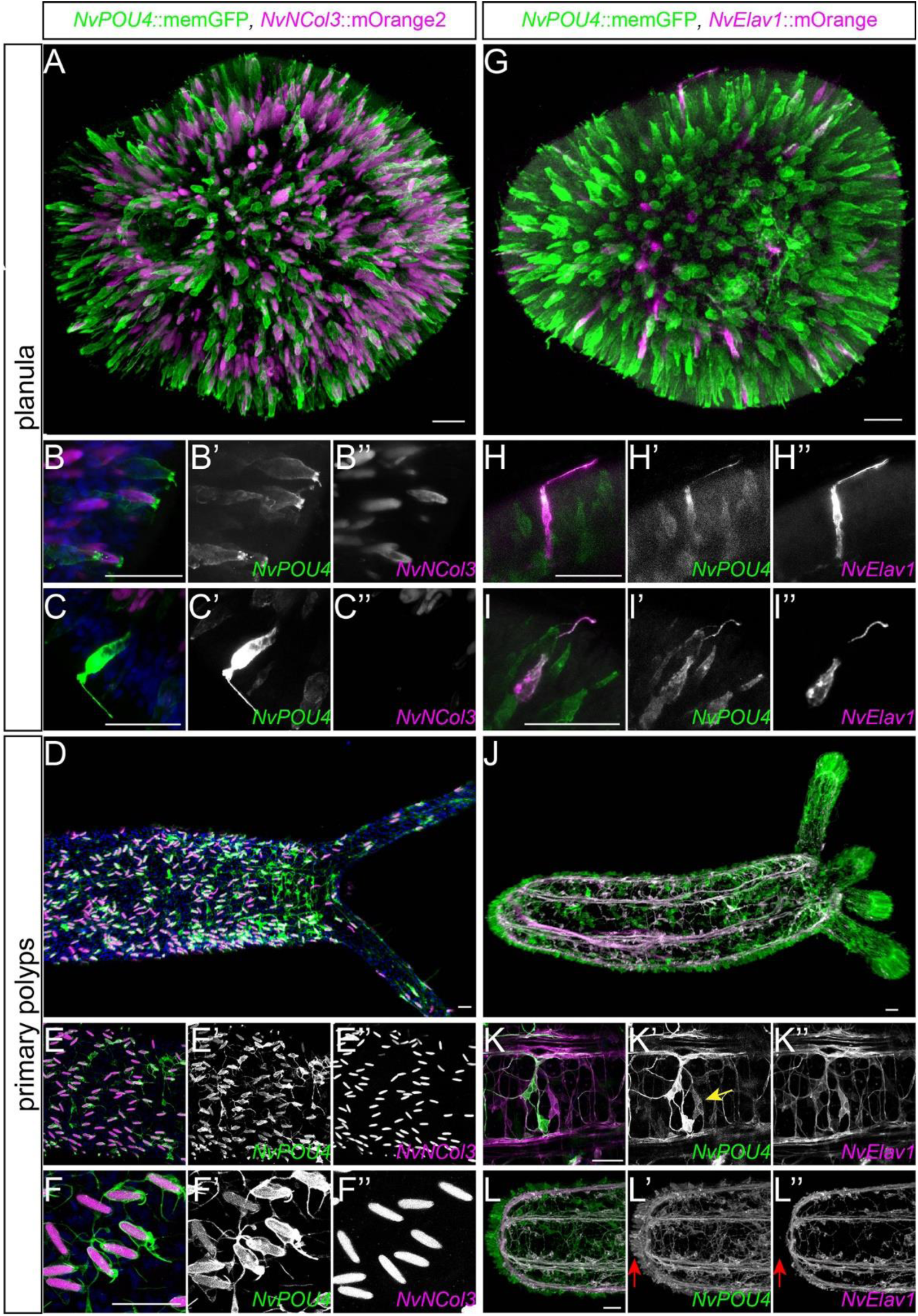
*NvPOU4*::memGFP identifies neural cell types. (A-F) Double transgenic animals with *NvPOU4*::memGFP (green) and *NvNCol3*::mOrange2 (magenta) and DAPI in blue. Lateral view with aboral pole to the left. (A-C) planula stage. (E-F) live images at primary polyp stage. (A,B,D,E,F) from gastrula to primary polyp stage the *NvNCol3*::mOrange2^+^ capsules are surrounded by the memGFP suggesting that *NvPOU4*::memGFP identifies cnidocytes. (C) Some *NvPOU4*::memGFP^+^ cells do not contain a *NvNCol3*::mOrange2^+^ capsules, suggesting that the *NvPOU4*::memGFP line identifies cnidocytes and other cell types. In (B-C and E-F), *NvPOU4* and *NvNCol3* stand for *NvPOU4*::memGFP and *NvNCol3*::mOrange2, respectively. (A, D) are projections of stacks, (B-C’’ and E-F’’) are single confocal sections. (G-L) Double transgenic animals with *NvPOU4*::memGFP (green) and *NvElav1*::mOrange (magenta). Lateral views with aboral pole to the left. (G-I) At planula stage, some ectodermal cells are positive for both reporter proteins, these cells have an apical cilium. (J-L) At primary polyp stage the endodermal nerve net expresses both fluorescent proteins. Arrow in (K’) indicates an endodermal cell with ganglion cell-like morphology. (L) The cnidocytes (red arrows in (L-L’) appear to be the only cells that do not co-express both transgenes. In (H-I and K-L), *NvPOU4* and *NvNCol3* stand for *NvPOU4*::memGFP and *NvNCol3*::mOrange2, respectively. (G, J) are projections of stacks, (H-I’’ and K-L’’) are single confocal sections. Scale bars represent 20 μm.

To gain further insight into the nature of these other *NvPOU4* expressing cells, we generated *NvPOU4*::memGFP, *NvElav1*::mOrange double transgenic animals (Figure 4G-L). The *NvElav1*::mOrange transgenic line labels a subset of sensory and ganglion cells but not cnidocytes (Nakanishi et al., 2012). From early planula stage, we could note ectodermal cells that expressed both fluorescent proteins, suggesting that a subset of sensory cells expresses *NvPOU4* (Figure 4 H, I). The co-expression of the two transgenes is more prominent at primary polyp stages, with much of the *NvElav1*::mOrange-positive endodermal nerve net also expressing the memGFP protein (Figure 4J-L), including cells with the morphology of ganglion cells (arrow in Fig. 4K’). We also generated double transgenics of *NvPOU4*::memGFP with the *NvFoxQ2d*::mOrange line, in which a small subpopulation of ectodermal sensory cells is labelled (Busengdal and Rentzsch, 2017). We did not observe co-expression of *NvPOU4*::memGFP with *NvFoxQ2d*::mOrange (data not shown).

Thus, the *NvPOU4* transgenic reporter line labels cnidocytes and a subset of sensory and ganglion cell types, and transgene expression is maintained at the polyp stage.

### *NvPOU4* is required for neural differentiation

To identify the function of *NvPOU4*, we generated a mutant line using the CRISPR/Cas9 system. We targeted the beginning of the POU domain with a sgRNA (Figure 5A). Genotyping of F1 animals derived from one founder polyp revealed a prevalent deletion of 31bp, causing a frame shift and a premature stop codon that truncates the encoded protein before the DNA binding domain (Figure 5A and S4). We denote this allele as *NvPOU4^1^* and refer to it in the text as *NvPOU4^-^*. We collected F1 animals with this 31bp deletion and then crossed them to obtain 25% homozygous F2 mutants. Light microscopic observation of the F2 animals showed that 26% of them lacked elongated cnidocyst capsules at primary polyp stage (n=97 animals). We extracted DNA and sequenced the *NvPOU4* gene of those animals and of their siblings that possessed cnidocysts (Figure 5B, C). This showed that 90% of the animals without cnidocysts were *NvPOU4^-/-^,* whereas all sibling controls with cnidocysts were *NvPOU4^+/+^*or *NvPOU4^+/-^*. This confirmed that *NvPOU4* homozygous mutants lack cnidocysts. These polyps were unable to catch prey and did not survive beyond the primary polyp stage. To better characterize the cnidocyte phenotype, we performed stainings to distinguish developing from mature cnidocytes. An antibody against NvNCol3 detects the cnidocysts throughout their development, but does not detect the fully mature cnidocyst capsules (with very few exceptions) (Babonis and Martindale, 2017; Zenkert et al., 2011). The matrix of the mature cnidocysts is specifically labelled by incubation with a high concentration of 4′,6-diamidino-2-phenylindole (DAPI) in the presence of EDTA (Szczepanek et al., 2002). We noticed that *NvPOU4^-/-^* polyps do have patches of NvNCol3 staining but lack any of the well-defined, elongated capsules that form during cnidocyte differentiation. Such capsules were neither visible by staining with the NvNCol3 antibody nor by the high concentration of DAPI (Figure 5D-K). We confirmed this observation by generating *NvPOU4^-/-^*, *NvNCol3*::mOrange2 animals; in these animals NvNCol3 is expressed but mature capsules fail to differentiate (Figure S5). The diffuse nature of NvNCol3 staining in the mutants made quantification of the stained cells difficult, thus we cannot exclude an effect on the number of NvCol3 expressing cells. These experiments suggest, however, that *NvPOU4* is primarily required for the terminal differentiation of cnidocytes in *Nematostella*.

**Fig. 5.**
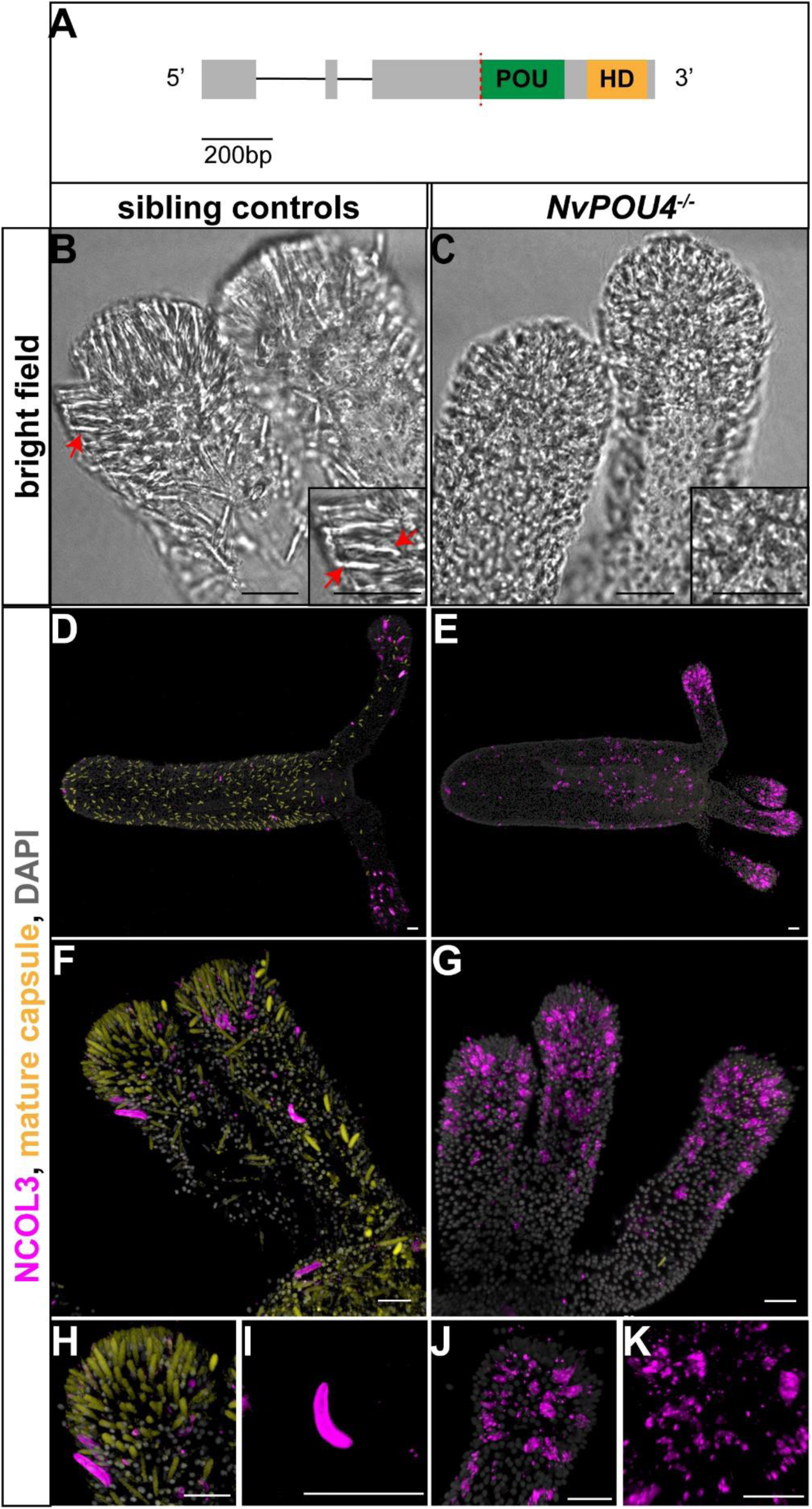
The loss of *NvPOU4* prevents cnidocyte differentiation. (A) Schematic of the CRISPR/Cas9 targeting strategy. Exons are in grey boxes, the POU domain is shown as a green box and the homeodomain (HD) as a yellow box. The sgRNA targets the start of the POU domain (red dashed line) and generated a deletion of 31bp causing a frame shift and the appearance of a premature STOP codon. (B, C) Bright field picture of the tentacle tips of (B) primary polyps control (*NvPOU4*^+/+^ and *NvPOU4*^+/-^) vs (C) *NvPOU4*^-/-^. A red arrow highlights an elongated cnidocyst. (D-K) antibody staining of NvNCol3 (magenta), mature capsules (yellow) and nuclei (grey) in controls (D, F, H, I) vs *NvPOU4*^-/-^ (E, G, J, K). *NvPOU4*^-/-^ animals lack elongated mature capsules. They still show NvNCol3 antibody staining, suggesting that cnidocytes are specified but do not differentiate properly, highlighted by the shape of the NvNCol3-positive capsule (I and K). (D-K) are projections of stacks of confocal sections. Scale bars represent 20 μm

We next studied the role of *NvPOU4* during the development of other neural cell types (ganglion and sensory cells) identified by the *NvPOU4*::memGFP reporter line. To do so, we generated *NvPOU4^-/-^, NvElav1*::mOrange animals by crossing *NvPOU4^+/-^* to *NvPOU4^+/-^*, *NvElav1*::mOrange polyps. We collected primary polyps with *NvElav1*::mOrange expression and inspected them for the cnidocyte phenotype. Despite a lack of mature cnidocytes, these putative *NvPOU4^-/-^, NvElav1*::mOrange animals displayed no gross aberration of their *NvElav1^+^* nervous system (Figure 6A-D). For quantification, we randomly imaged *NvElav1*::mOrange^+^ polyps, counted the number of mOrange^+^ cells in a 100 μm × 100 μm square in the body column and determined the presence or absence of mature cnidocytes by light microscopy a posteriori. We did not find a statistically significant difference in the number of mOrange^+^ cells between polyps with and without mature capsules, respectively (Figure S6). This suggests that *NvPOU4* does not have a major role in the specification or gross morphological development of the *NvElav1^+^* neurons.

**Fig. 6.**
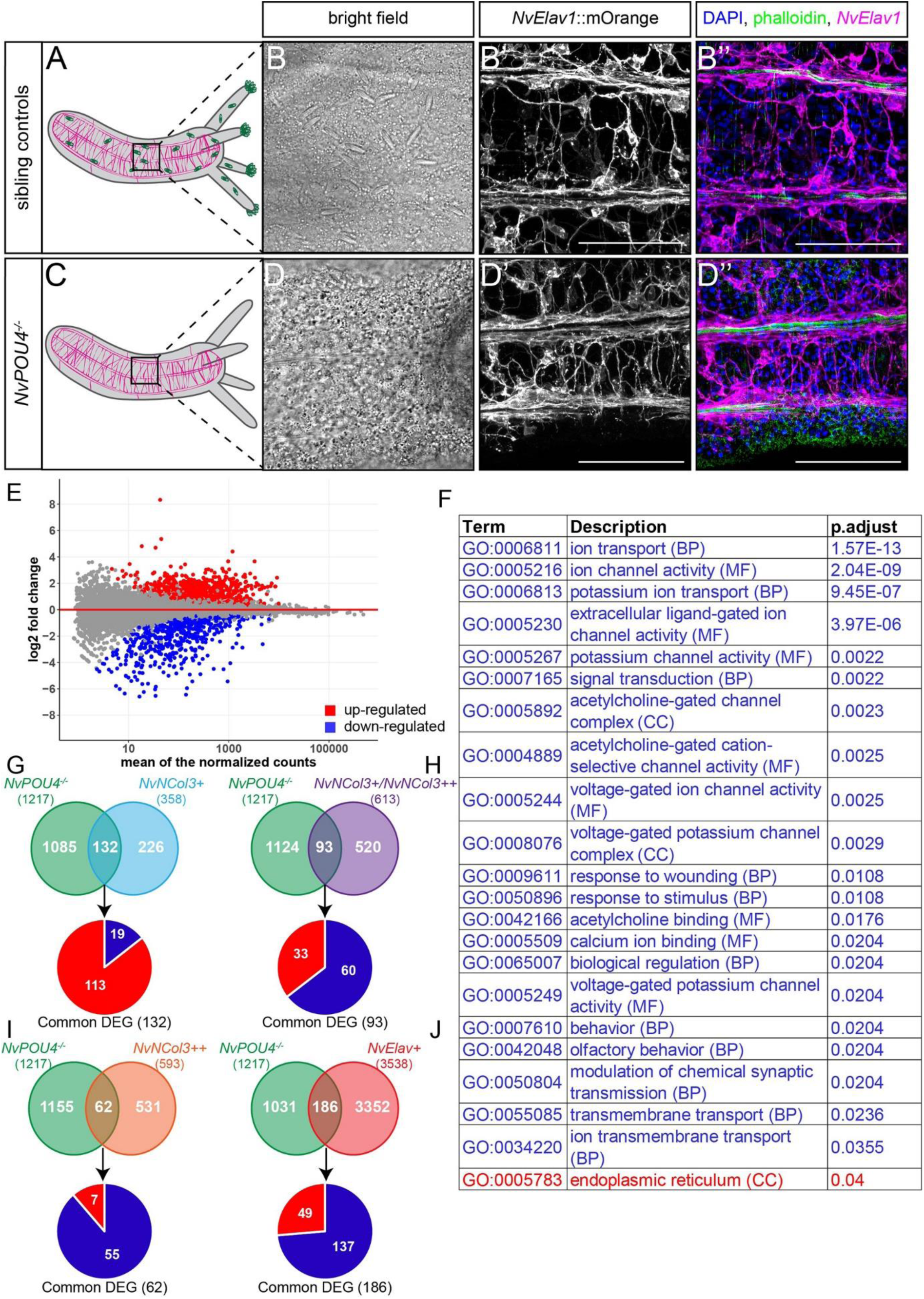
Loss of *NvPOU4* affects the transcriptomes of *NvNCol3* and *NvElav1* expressing neural cells. (A-D) *NvPOU4^-/-^* animals were distinguished based on the absence of cnidocyte capsules. (B’-D’’) Confocal images of anti dsRed antibody staining (detecting mOrange, shown in magenta) in *NvPOU4^-/-^, NvElav1::*mOrange^+/-^ polyps (10 dpf), phalloidin in green, DAPI in blue. Scale bars represent 50 μm. This experiment suggested that the *NvElav1*^+^ cells are specified properly in the absence of *NvPOU4.* (E) MA plot of the RNA sequencing comparing *NvPOU4^-/-^* animals with their sibling control (selection based on the cnidocyte phenotype, four biological replicates). In total 1217 genes were differentially expressed (p-adjusted <0.05, no minimal fold change), 641 were found up-regulated (red) and 576 were found down-regulated (blue). (F) GO term analysis of the *NvPOU4^-/-^* vs siblings (with p<0,05). GO terms overrepresented among downregulated genes in blue, those overrepresented among upregulated genes in red. (G-H) Comparison of the genes differentially expressed in *NvPOU4^-/-^* with the published *NvNCol3::*mOrange2 transcriptomes (Sunagar et al.2018). *NvNCol3^+^* represents the genes expressed in differentiating cnidocytes (358), *NvNcol3^++^* represents the genes expressed in fully differentiated cnidocytes (593) and *NvNcol3^+^/NvNcol3^++^*represents the genes that are expressed in both, differentiating and differentiated cnidocytes (613). (G) 85.6% (113/132) of the differentially expressed genes common to the *NvPOU4* mutant and the *NvNcol3^+^* transcriptomes are up-regulated in the *NvPOU4* mutants. (H) 64.5% (60/93) of the differentially expressed genes common to *NvPOU4* mutant and the *NvNcol3^+^/NvNcol3^++^* transcriptome are down-regulated in the *NvPOU4* mutants. (I) 85.6% (55/62) of the differentially expressed genes common to the *NvPOU4* mutant and the *NvNcol3^++^* transcriptome are down-regulated in the *NvPOU4* mutants. (J) Comparison of the genes differentially expressed in *NvPOU4^-/-^* with the *NvElav1*::mOrange transcriptome. 73.7% (137/186) of the differentially expressed genes common to *NvPOU4* mutant and the *NvElav1* transcriptome are down-regulated in the *NvPOU4* mutants.

### Loss of *NvPOU4* affects the transcriptomes of *NvNCol3* and *NvElav1*-expressing neural cells

To characterize the function of the *NvPOU4^-^* in more detail, we decided to analyze transcriptional changes using RNA sequencing. We compared cnidocyst-lacking *NvPOU4^-/-^* animals at primary polyp stage and compared them to their cnidocyst-containing siblings (consisting of *NvPOU4^+/+^* and *NvPOU4^+/-^* animals). RNA sequencing of four biological replicates confirmed that *NvPOU4^-/-^* animals only generate transcripts of this gene with the 31bp deletion (Figure S7). In total, 1217 genes were differentially regulated (p-adjusted value <0.05, no threshold for fold-change), with 576 being down- and 641 being up-regulated in the *NvPOU4^-/-^* polyps (Figure 6E and Table S1). An analysis of Gene Ontology (GO) terms identified 21 terms that are overrepresented among the downregulated genes, with “ion channel activity”, “extracellular ligand-gated ion channel activity”, “potassium channel activity”, “acetylcholine binding”, “calcium ion binding” and “voltage-gated potassium channel activity” being overrepresented in the GO domain “Molecular function” (Figure 6F). The only term that is overrepresented among the up-regulated genes is “endoplasmic reticulum” in the GO domain “Cellular component”. While this is consistent with a role for *NvPOU4* in nervous system development, the low proportion of *Nematostella* genes that are associated with a GO term (39.4% of all differentially expressed genes) limits the power of the analysis.

Our analysis of double transgenic animals has shown that separate subsets of *NvPOU4*-expressing cells give rise to *NvNCol3*- and *NvElav1*-expressing cells (Figure 4). We therefore used the transcriptomes of *NvNCol3*::mOrange^+^ (Sunagar et al., 2018) and *NvElav1*::mOrange^+^ cells enriched by FACS, to assign genes differentially regulated in *NvPOU4* mutants to these two different populations of neural cells. For the *NvNCol3*::mOrange^+^ cells, different levels of fluorescence combined with microscopic examination have previously been used to characterize two populations of these cells and to generate transcriptomes of them. One of the cell populations is enriched for differentiating cnidocytes (mOrange positive) and the other one consists of mature cnidocytes (mOrange super-positive, with higher fluorescence) (Sunagar et al., 2018). Among the genes that were differentially expressed in *NvPOU4* mutants, we identified 287 genes that were in common with those reported in the *NvNCol3*::mOrange transcriptomes, with 132 genes being differentially expressed only in the positive cells, 62 only in the super-positive cells and 93 in both groups of cells (Figure 6G-I and Table S2). Interestingly, we noticed a striking difference in the proportion of up- and downregulated genes in these three groups of cells. Of the 132 differentially expressed genes (DEGs) present only in the mOrange-positive, differentiating cells, 113 (85.6%) were up- and 19 (14.4%) were down-regulated (Figure 6G) in the *NvPOU4* mutants. In contrast, of the 62 DEGs only present in the mOrange super-positive, more mature cells, 55 (88.7%) were down-, and only 7 (11.3%) were upregulated (Figure 6I). Of the 93 DEGs found in both groups of cells, 60 (64.5%) were down- and 33 (35.5%) were upregulated (Figure 6H). This suggests that mutation of *NvPOU4* reduces the expression of genes involved in the terminal differentiation of cnidocytes and increases the expression of genes involved in earlier steps of their development.

We next generated transcriptomes of *NvElav1*::mOrange-positive and negative cells at primary polyp stage and identified 3538 genes with significantly higher expression level in *NvElav1*::mOrange-positive cells. Of these genes, a total of 186 were differentially regulated in *NvPOU4* mutants and among them, 137 (73.7%) were downregulated, but only 49 (26.3%) were upregulated in the mutants (Figure 6J). Thus, similar to the situation in the *NvNCol3*::mOrange super-positive cnidocytes, *NvPOU4* appears to function mainly as a positive regulator of genes expressed in *NvElav1*::mOrange positive cells. The broad expression in *NvElav1*::mOrange^+^ neurons suggests that *NvPOU4* functions in the differentiation of different types of neurons. To support this hypothesis, we selected two genes that are downregulated in the *NvPOU4* mutants and are upregulated in *NvElav1*::mOrange expressing cells for double fluorescent in situ hybridization with *NvPOU4*. We found *Nve22966* (a putative ionotropic glutamate receptor) to be co-expressed with *NvPOU4* mainly in endodermal cells (Figure 7A,B), whereas the co-expression of *NvPOU4* and *Nve21438* (a putative GABAA receptor subunit) is most prominent in ectodermal cells (Figure 7C, D). This indicates that *NvPOU4* indeed contributes to the differentiation of different subpopulations of *NvElav1*::mOrange expressing neurons.

**Fig. 7.**
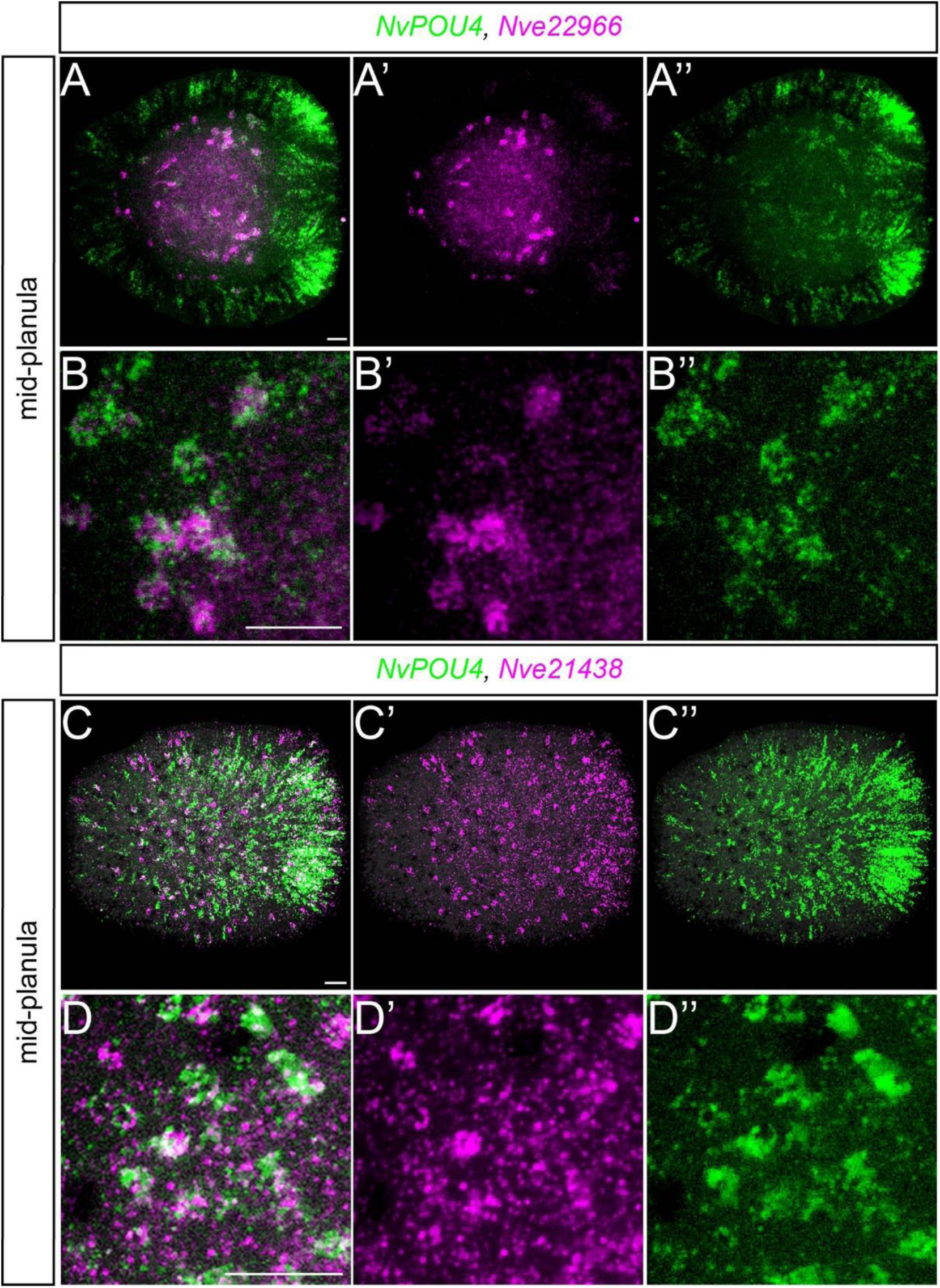
Co-expression of *NvPOU4* with glutamate and GABA_A_ receptor genes in different populations of cells. (A-D) Lateral views of double fluorescence *in situ* hybridization with probes indicated on the top and the developmental stage on the left side. (A-B) *NvPOU4* is labelled in green and *Nve22966* (a putative glutamate receptor) in magenta, co-expression is visible in an endodermal population of cells. (C-D) *NvPOU4* is labelled in green and *Nve21438* (a putative GABAA receptor subunit) in magenta, co-expression is visible in an ectodermal population of cells. This suggests *NvPOU4* contributes to the differentiation of different subpopulations of *NvElav1*::mOrange^+^ cells. Lateral views with the aboral pole to the left. (A-A’’ and C-C’’) are projections of stacks of confocal sections, (B-B’’ and D-D’’) are single confocal sections. Scale bars represent 20 μm.

Taken together, our transcriptome analyses show that *NvPOU4* is mainly required for the expression of genes that are expressed at late stages of cnidocyte and neuron differentiation. This is consistent with the morphological observations of the *NvPOU4* mutants in the background of the *NvNCol3*::mOrange2 and *NvElav1*::mOrange transgenic lines, and suggests that *NvPOU4* regulates the terminal differentiation of neural cells.

## DISCUSSION

In this report we have shown that *NvPOU4* is expressed in postmitotic cells that are derived from *NvSoxB(2)* expressing neural progenitor cells and give rise to cnidocytes, sensory cells and ganglion cells. Despite its expression from blastula stage on, mutation of *NvPOU4* does not prevent the initial specification of these cells. Instead, our data suggest that the main function of *POU4* in *Nematostella vectensis* is the regulation of the terminal differentiation of neural cells.

Cnidocyte precursors in *NvPOU4* mutants still produce NvNcol3 protein. They fail, however, to assemble the elongated cnidocysts characteristic of mature cnidocytes. This morphological observation suggested that in these cells, *NvPOU4* mainly regulates the transcription of genes that encode factors required for the final steps of cnidocyte differentiation. The availability of separate transcriptomes enriched for cnidocytes at earlier stages (at or before the beginning of cnidocyst formation) and at later stages (containing elongated cnidocysts) of their development (Sunagar et al., 2018) allowed us to analyze the requirement for *NvPOU4* in this process in more detail. The preponderance of downregulation among genes that are specifically enriched in late-stage cnidocytes matched the observed lack of mature cnidocysts. In contrast, we did not expect that many of the genes that are specifically expressed in early-stage cnidocytes would be upregulated in *NvPOU4* mutants. This included the *NvNCol3* gene, *NvPaxA* (a transcription factor that positively regulates *NvNCol3* expression, (Babonis and Martindale, 2017) and *NvPOU4* itself. A possible explanation is that the failure to produce functional cnidocytes leads to a “compensatory” response that increases the number of cells entering the cnidocyte differentiation pathway. An alternative, but not mutually exclusive possibility is that *NvPOU4* is required for the downregulation of genes that are temporarily expressed at an earlier stage of cnidocyte differentiation, resulting in prolonged expression of these genes in *NvPOU4* mutants.

For the *NvElav1*::mOrange-expressing neurons, we currently cannot separate cells at different stages of their differentiation. At the primary polyp stage, all or almost all *NvElav1*::mOrange positive neurons possess neurites and are thus either at a late stage of their development or terminally differentiated (Nakanishi et al., 2012). In homozygous *NvPOU4* mutants, there is no significant reduction in the number of *NvElav1*::mOrange expressing neurons and the neurons extend neurites that do not show gross morphological alterations or obviously aberrant projection patterns (Figure 6B’, D’). These observations suggest that in the *NvElav1^+^* endodermal neurons *NvPOU4* mainly functions in terminal differentiation, regulating for example the repertoire of neurotransmitter receptors. In line with such a function, the expression levels of genes encoding glutamate, acetylcholine and GABA receptors are reduced in *NvPOU4* mutants. We note, however, that *NvPOU4* may have other or additional roles in subpopulations of *NvElav1*-expressing neurons.

Of the 1217 genes differentially expressed in *NvPOU4* mutants, only 287 are upregulated in the cnidocyte transcriptomes and 186 in the *NvElav1*::mOrange transcriptome. This is a surprising observation since the double transgenic lines suggest that the majority of *NvPOU4*::memGFP expressing cells are included in the *NvNCol3*::mOrange or *NvElav1*::memOrange-positive cells. A possible explanation is the difference in the age of the polyps used for the isolation of cnidocytes (3-4 months, (Sunagar et al., 2018) and the *NvPOU4* mutants (14 days). The proportion of different types of cnidocytes has been shown to differ between primary and adult polyps (Zenkert et al., 2011) and the cnidocyte transcriptomes may therefore lack genes that are expressed predominantly at earlier stages. Similarly, only cnidocytes from the tentacles were used for generating the cnidocyte transcriptomes, whereas *NvPOU4* is expressed in both tentacle and body column cnidocytes and these regions have been shown to differ in the composition of cnidocyte types (Zenkert et al., 2011). Genes that are expressed in cnidocyte types that are more common in the body column (e.g. basitrichous haplonemas) may therefore be underrepresented in the cnidocyte transcriptomes. It will be interesting for future studies to understand whether additional populations of *NvPOU4*-expressing cells exist outside the *NvNCol3* and *NvElav1*^+^ cells.

After functioning in neural development, POU4 genes have been shown to be required for the survival of several classes of neurons in *C.elegans* and in the mouse habenula (Serrano-Saiz et al., 2018). Deletion of *POU4* in these terminally differentiated neurons results in the loss of their neurotransmitter identity and their elimination by apoptosis (Serrano-Saiz et al., 2018), suggesting an evolutionarily conserved, post-developmental role for POU4 genes. Whether *NvPOU4* has a comparable role in *Nematostella* is currently not clear. A recent single cell RNA sequencing study showed that *NvPOU4* is expressed in neural cells in adult animals (Sebe-Pedros et al., 2018), allowing for a role in maintaining neural identity. We observed homozygous mutants until 20 days post fertilization (they become primary polyps after 6-7 days), but did not detect alterations in the number or morphology of *NvElav1*::mOrange positive neurons (data not shown). Due to the lack of cnidocytes, *NvPOU4* mutants are unable to catch prey and are thus not viable, which prevents long-term observations. Determining whether *NvPOU4* has a role in the maintenance of the identity of neurons will require the development of methods for conditional gene inactivation in *Nematostella*.

The *C. elegans* POU4 gene *unc-86* is a prime example of a terminal selector gene and POU4 genes in other species have comparable functions in the terminal differentiation of neural cell types (Ayer and Carlson, 1991; Certel et al., 2000; Clyne et al., 1999; Duggan et al., 1998; Erkman et al., 2000; Gan et al., 1996; Gordon and Hobert, 2015; Huang et al., 2001; Huang et al., 1999; Serrano-Saiz et al., 2013; Sze et al., 2002). Terminal selectors act in a combinatorial manner to regulate terminal effector genes and determine cellular identity (Hobert, 2016). This allows individual transcription factors to act as terminal selectors in different types of neurons, for example by cooperative binding to regulatory elements together with other transcription factors (Cho et al., 2014; Duggan et al., 1998; Wolfram et al., 2014; Xue et al., 1992). In *Nematostella*, single-cell RNA sequencing has revealed several *NvPOU4* expressing “metacells” with overall neuron-like transcriptional profiles (Sebe-Pedros et al., 2018). Individual *NvPOU4*^+^ metacells (likely representing different neural cell types) express different combinations of other transcription factors (Sebe-Pedros et al., 2018), some of which may function together with *NvPOU4* in regulating the terminal differentiation of these cells. In line with this scenario, each of the *NvPOU4*^+^ metacells expresses at least one transcription factor that has not been detected in any other *NvPOU4*^+^ metacell (Table S3). While the physical and functional interaction with other transcription factors and with regulatory elements of target genes remains to be explored in future work, the morphological and molecular analyses presented here support the hypothesis that *NvPOU4* acts as a terminal selector for different neural cell types in *Nematostella*.

The advent of single-cell sequencing technologies has led to the elaboration of concepts for the evolutionary diversification of cell types. Terminal selector genes are central to such concepts as they are part of so-called “core regulatory complexes” (CoRCs) which regulate the cellular features that distinguish different cell types (Arendt et al., 2016). It has been hypothesized that evolutionary changes occur more slowly in core regulatory complexes than in terminal effector genes, which would make them more informative for inferring evolutionary relationships between cell types (Arendt et al., 2019; Arendt et al., 2016). Our data show that more than 600 million years after the divergence of the cnidarian and bilaterian lineages (dos Reis et al., 2015), POU4 genes function in the terminal differentiation of neural cells in cnidarians, as they do in bilaterians. *NvPOU4* regulates the terminal differentiation of strikingly different types of neural cells in *Nematostella*, the more “typical” *NvElav1*^+^ neurons and the highly derived, taxon-specific cnidocytes. This is likely due to cell type-specific combinatorial regulation together with other transcription factors and attempts to homologize POU4-expressing neural cell types will require a more detailed understanding of such combinatorial regulation. Nevertheless, we propose that the function of *NvPOU4* is derived from an ancestral function of POU4 genes as regulators of terminal neural differentiation. We cannot exclude, however, that in some cases POU4 genes have been co-opted into comparable roles in different cell types.

In summary, the observation that *NvPOU4* functions in the terminal differentiation of neural cells in *Nematostella* supports the hypothesis that the regulation of neurogenesis by conserved terminal selector genes is an ancient feature of nervous system development.

## STAR METHODS

### KEY RESOURCES TABLE

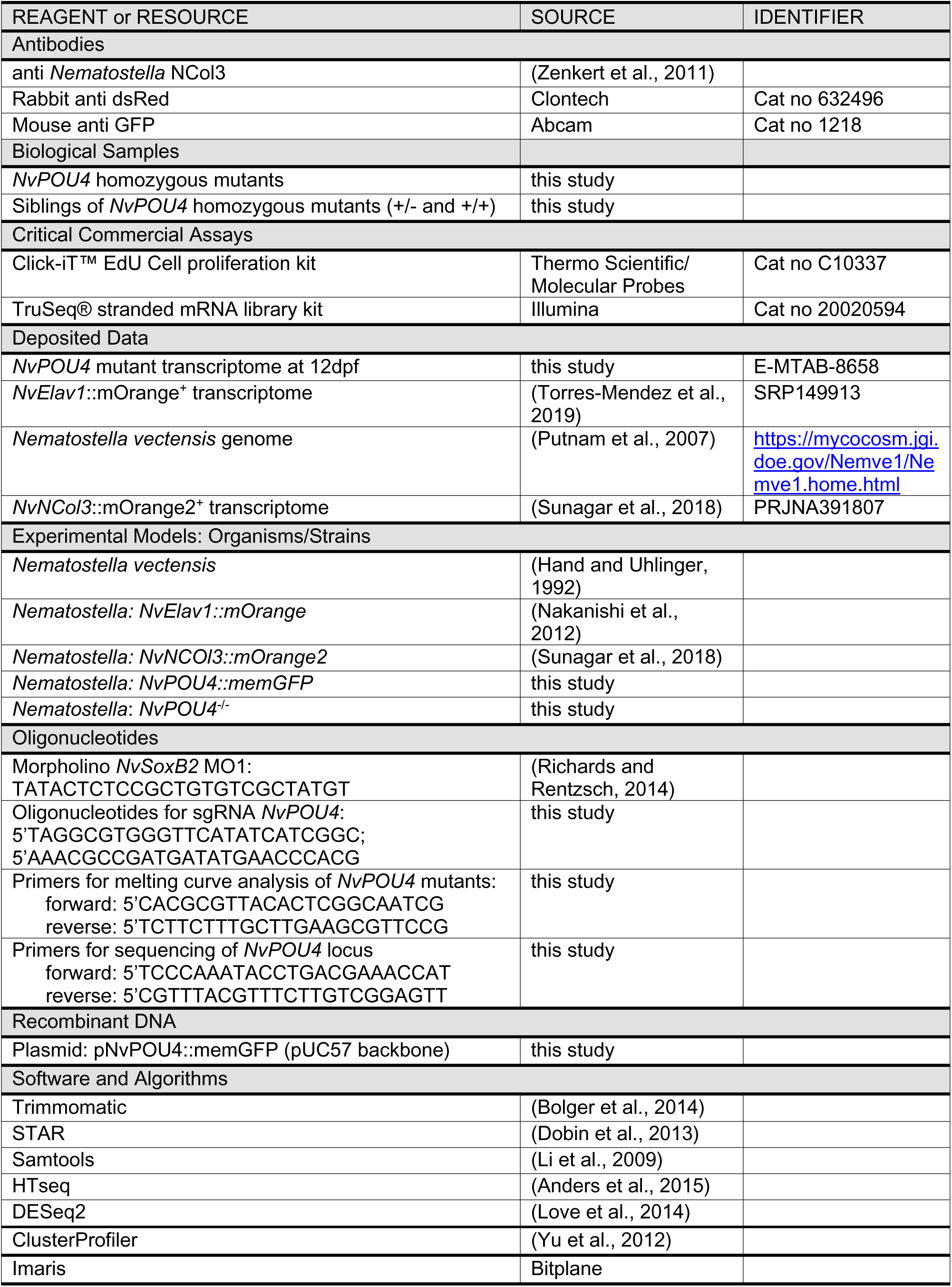

### Nematostella culture

Adult animals were maintained at 18 °C in 1/3 filtered seawater (=*Nematostella* medium, NM). Spawning induction was performed by light and temperature shift as described in (Fritzenwanker and Technau, 2002). Incubation of the fertilized egg packages with a 3% cysteine/NM removed the jelly. Embryos were then raised at 21 °C and fixed at 12 hours post fertilization (hpf; early blastula), 16hpf (blastula), 20hpf (early gastrula), 24hpf (gastrula), 30hpf (late gastrula), 48hpf (early planula), 72hpf (planula), 4dpf (late planula); 5dpf (tentacle bud); 7dpf (early primary polyp), 12dpf (late primary polyp).

### Morpholino injection

*NvSoxB(2)* MO1 is described in (Richards and Rentzsch, 2014). Experiments were conducted with four biological replicates, with embryos derived from four independent spawnings.

### Generation of transgenic lines

The *NvPOU4*::memGFP transgenic reporter line was generated by meganuclease-mediated transgenesis as described by (Renfer and Technau, 2017). The genomic coordinates for the 4.7 kb regulatory region are 1063816-1068603 on scaffold 16 (http://genome.jgi.doe.gov/Nemve1/Nemve1.home.html, accessed 15 April 2019). This fragment was inserted in front of a codon optimized GFP via the HiFi DNA Assembly kit (NEB) with the addition of a membrane-tethering CAAX domain at the C-terminus to visualize the morphology of the cells expressing the reporter protein. memGFP was detected with an anti-GFP antibody (mouse, abcam1218, 1:200). The *NvSoxB(2)*::mOrange line has been described in (Richards and Rentzsch, 2014), *NvElav1*::mOrange in (Nakanishi et al., 2012), *NvNCol3*::mOrange in (Sunagar et al., 2018) and *NvFoxQ2d*::mOrange in (Busengdal and Rentzsch, 2017). An overview of the crosses are provided in Table S4.

### *Cloning of NvPOU4, in situ* hybridization, EdU labelling and immunohistochemistry

The *NvPOU4* sequence is derived from gene model Nve5471, retrieved from https://figshare.com/articles/Nematostella_vectensis_transcriptome_and_gene_models_v2_0/807696. Fluorescent and colorometric *in situ* hybridizations were performed as described in the Supplementary material in (Richards and Rentzsch, 2014). Samples were imaged on either a Nikon Eclipse E800 compound microscope with a Nikon Digital Sight DSU3 camera or on a Leica SP5 confocal microscope.

The following primary antibodies were used: to detect *NvPOU4*::memGFP, anti-GFP (mouse, abcam1218, 1:200); to detect mOrange, anti dsRed (rabbit, Clontech 632496, 1:100); anti-NCol3 (Zenker et al.2011); mature cnidocytes were labelled with DAPI/EDTA as described in (Babonis and Martindale, 2017; Szczepanek et al., 2002).

EdU labelling was done as 30min pulses followed by fixation as described in (Richards and Rentzsch, 2014), using Click-it EdU Alexa fluor 488 kit (Molecular probes C10337). For counting *NvPOU4^+^* and EdU^+^ cells a 100 μm × 100 μm sampling area was defined in the mid-lateral region of the ectoderm at blastula stage. All the nuclei from this region were scanned via confocal microscopy.

### CRISPR-Cas9 mediated mutagenesis and genotyping of embryos

Using published methods (Ikmi et al., 2014; Kraus et al., 2016) sgRNA were synthesized i*n vitro* via the Megashortscript T7 kit (Life technologies) using the following oligos: 5’TAGGCGTGGGTTCATATCATCGGC, 5’AAACGCCGATGATATGAACCCACG

The reaction mixture (500 ng/μl Cas9 enzyme and 150 ng/μl of the sgRNA) was incubated at 37°C for 15min prior to injection.

Genomic DNA from embryos or aboral pieces of F1 polyps was extracted using a Tris/EDTA/proteinase K buffer. Mutant genotyping was first done via melt-curve analysis after PCR amplification of a 90bp region on a BioRad CFX96 RealRime PCR machine. Mutations were confirmed by sequencing a 500bp region around the mutation. Primers used are listed in the Key Resources Table.

### Generation of transcriptomes from *NvPOU4* mutants and siblings

The presence/absence of cnidocysts was used for sorting animals at primary polyp stage (12dpf) into sibling control (*NvPOU4^+/+^* and *NvPOU4^+/-^)* and mutants (*NvPOU4^-/-^*). Twenty primary polyps were pooled for each biological condition and the total RNA was extracted using the Direct-zol RNA MicroPrep kit (Zymo Research). Experiments were conducted with four biological replicates, with embryos derived from four independent spawnings. Sequencing libraries were generated with the TruSeq® stranded mRNA library prep kit (Illumina), 75bp single read sequencing was performed on a NextSeq500 machine (Illumina).

### Cell type specific transcriptomes

*NvElav1*::mOrange-positive cells were enriched by FACS and RNA was extracted as described previously (Torres-Mendez et al., 2019). cDNA was prepared from 400pg of total RNA using the Smart-Seq 2 method with 16 pre-amplification PCR cycles, as described by (Picelli et al., 2014). NGS libraries were prepared using the home-made tagmentation-based method as described by (Hennig et al., 2018). Briefly, 125 ng of cDNA was tagmented using home-made Tn5 loaded with annealed linker oligonucleotides for 3 minutes at 55C. Reaction was inactivated by adding 1.25ul of 0.2% SDS and incubation for 5 minutes at room temperature. Indexing and amplification was done using the KAPA HiFi HotStart PCR kit (Sigma-Aldrich) with Index oligonucleotides (sequences were adapted from Illumina). Four biological replicates of mOrange-positive and -negative cells, respectively, were used for 75bp single read sequencing on a NextSeq500 machine (Illumina).

The generation *NvNCol3*::mOrange2 transcriptomes is described in (Sunagar et al., 2018).

### Transcriptome analyses

The raw fastq files were initially quality checked and trimmed using Trimmomatic (Bolger et al., 2014). Following this they were aligned to the *N. vectensis* genome (https://mycocosm.jgi.doe.gov/Nemve1/Nemve1.home.html) using STAR (Dobin et al., 2013) in two-pass mode. Afterwards the produced BAM (Binary Alignment Maps) files were sorted and indexed with Samtools (Li et al., 2009) and then gene counting was carried out using HTSeq (Anders et al., 2015). Gene models were retrieved from (https://figshare.com/articles/Nematostella_vectensis_transcriptome_and_gene_models_v2_0/807696).

Differential gene expression testing and subsequent over-representation analysis was done in R with DESeq2 (Love et al., 2014) and clusterProfiler (Yu et al., 2012), respectively.

### Quantification of *NvElav1*::mOrange^+^ cells in *NvPOU4* mutants and siblings

*NvElav1*::mOrange positive cells (stained with anti dsRed antibody) were counted in an area 100µm long and located between two mesenteries. The quantification was done in animals from four different spawnings and with 5-10 animals per sample. The genotype was inferred a posteriori by the presence/absence of cnidocysts.

## ACKNOWLEDGEMENTS

We thank James Gahan and Marta Iglesias for generating the *NvElav1*::mOrange samples for RNA sequencing; Suat Özbek (COS Heidelberg) for the NvNCol3 antibody; James Gahan for critical reading of the manuscript; Eilen Myrvold and Lavina Jubek for excellent care of the *Nematostella* facility and Ivan Kouzel for help with data visualization. Clemens Döring provided advice on Imaris and Malalaniaina Rakotobe helped with initial cloning and in situ hybridizations. Sequencing of the *NvPOU4* mutant and sibling transcriptomes was performed at the Norwegian Sequencing Centre (Oslo), sequencing of the *NvElav1*::mOrange transcriptome at EMBL GeneCore (Heidelberg). Cell sorting was performed at the Flow Cytometry Core Facility, Department of Clinical Science, University of Bergen. The work was funded by the Sars Centre Core budget and by a grant from the Research Council of Norway and the University of Bergen (251185/F20) to FR; work in the Moran group was supported by grant 869/18 of the Israel Science Foundation to YM; KS was supported by funding from the DBT-IISc Partnership Program.

## Author Contributions

O.T. designed and performed the experimental work, analyzed the data, contributed to the conceptualization, generated the figures and wrote the manuscript. D.D. analyzed the transcriptome data. G.S.R. performed initial expression analysis of *NvPOU4* and provided supervision in the beginning of the project. K.S., Y.Y. C.-S. and Y.M. generated the *NvNCol3::mOrange2* transgenic line and shared it prior to publication. F.R. conceptualized and supervised the study and wrote the manuscript. All authors commented on the manuscript.

