## Supplemental Figures S1-7 for "*NvPOU4/Brain3* functions as a terminal selector gene in the nervous system of the cnidarian *Nematostella vectensis*"

### SUPPLEMENTAL INFORMATION

- **Supplementary figures and figure legends (Fig S1 – S7)**
- **Legends for Supplementary tables S1-S4**
- **Legends for Supplementary movies S1 – S10**

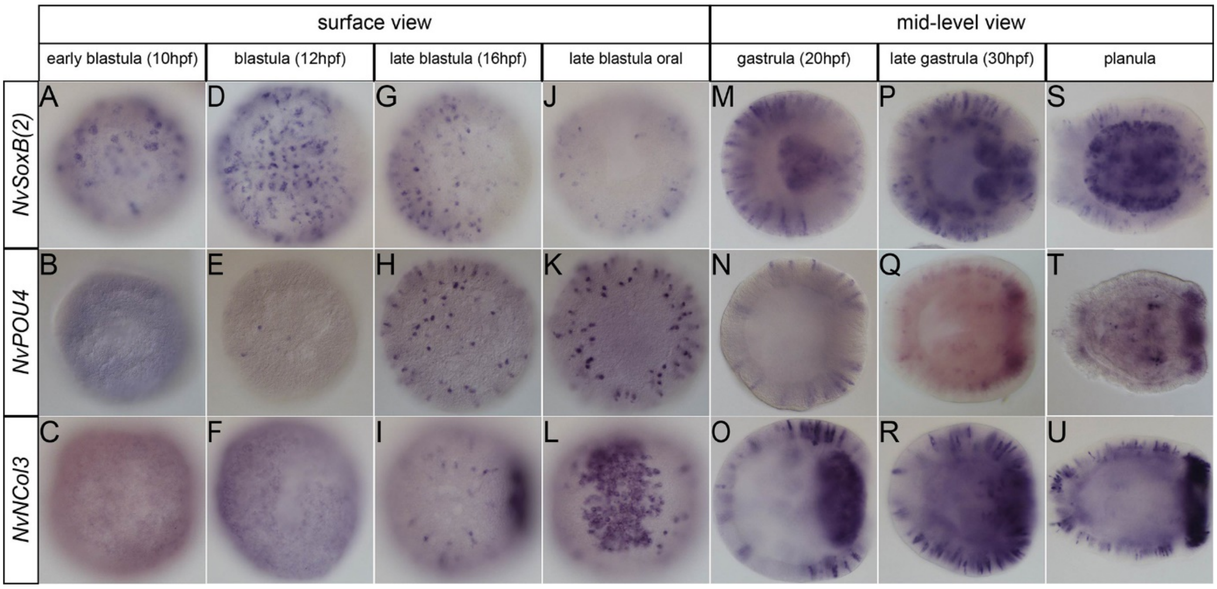


**Fig. S1 Related to Fig. 1 Relative expression of *NvSoxB(2)*, *NvPOU4* and *NvNcol3* from early blastula stage to late planula stage.** *In situ* hybridization with probes indicated on the left side and the developmental stage on top. (G-U) are lateral views with the aboral pole to the left, except for (J-L) which are oral views. (A-L) From Early blastula to late blastula, the pictures show the ectodermal surface view of the embryo. (M-U) From gastrula to planula stage, the pictures show the middle view of the embryo.

(A-C) *NvSoxB(2)* is the first of the three gene to be expressed at 10hpf, (D-F) followed at 12hpf by the first *NvPOU4* expressing cells. (G-I) *NvNcol3* start to be expressed around 16hpf. (J-L) *NvNcol3* is strongly expressed at the oral pole before gastrulation, which is not the case for *NvPOU4* and *NvSoxB(2)*. (M-U) From gastrula stage to planula stage the expression of *NvPOU4* and *NvNcol3* becomes more and more similar by being expressed in scattered single cell in the ectoderm and then in the future tentacle buds, whereas *NvSoxB(2)* is strongly expressed in the pharynx and in the ectoderm. All the embryos used for this experiment came from the same batch. Some images for *NvPOU4* are the same as in Fig 1.

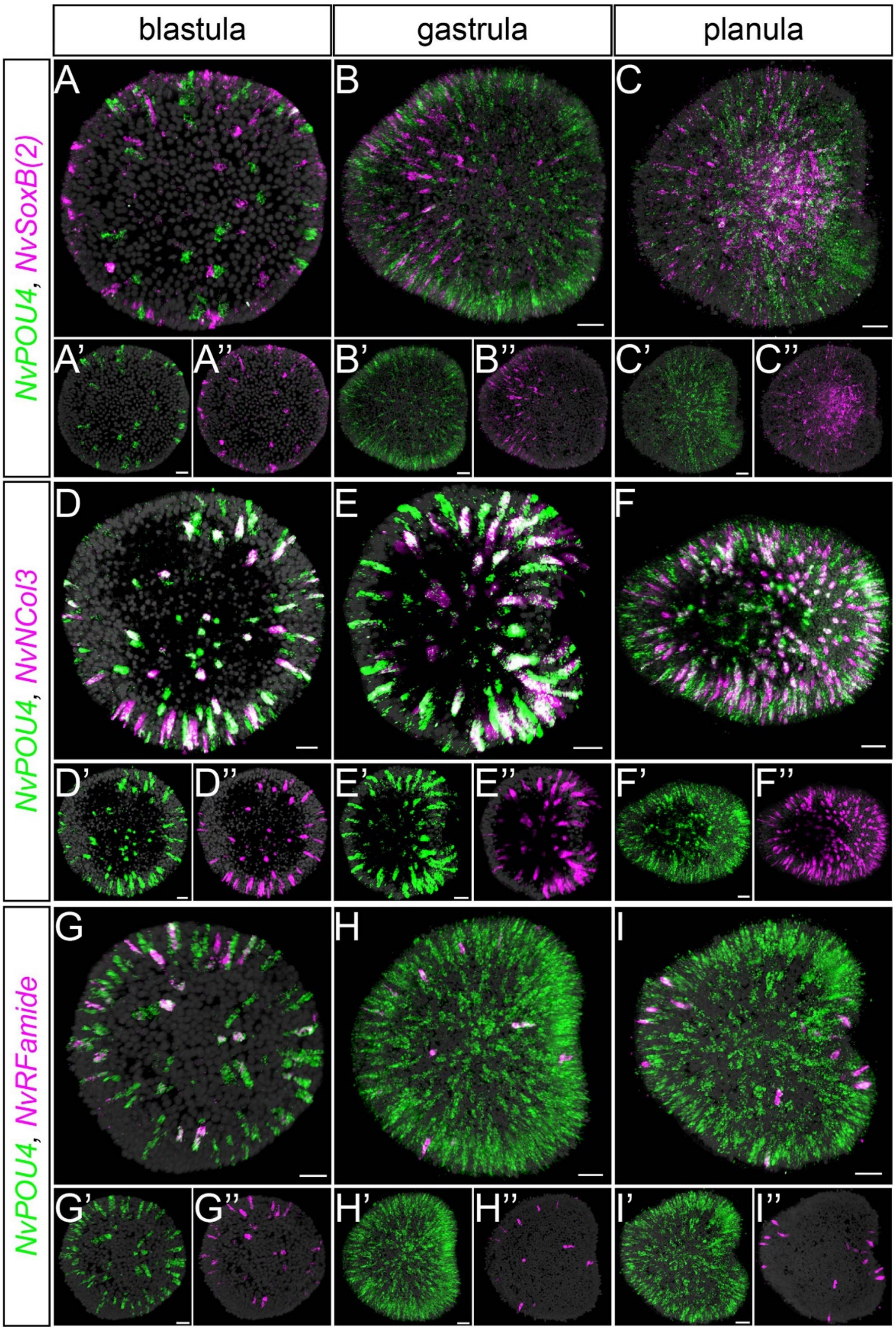


**Fig. S2 related to Fig. 2 *NvPOU4* is expressed in differentiating neural cells from blastula to planula stage.** (A-I) Double fluorescent *in situ* hybrization with probes indicated on the left side and the developmental stage on top, Lateral views, for gastrula and planula stages the aboral pole to the left. *NvPOU4* is shown in green, other probes in magenta: (A-C) *NvSoxB(2)*, (D-E) *NvNcol3* and (G-I) *NvRFamide*. All images are projections of stacks of confocal sections. Stacks for blastula, gastrula and planula stages are available as Supplementary movies 1-3, 4-6 and 7-9, respectively. Scale bars represent 20 µm.

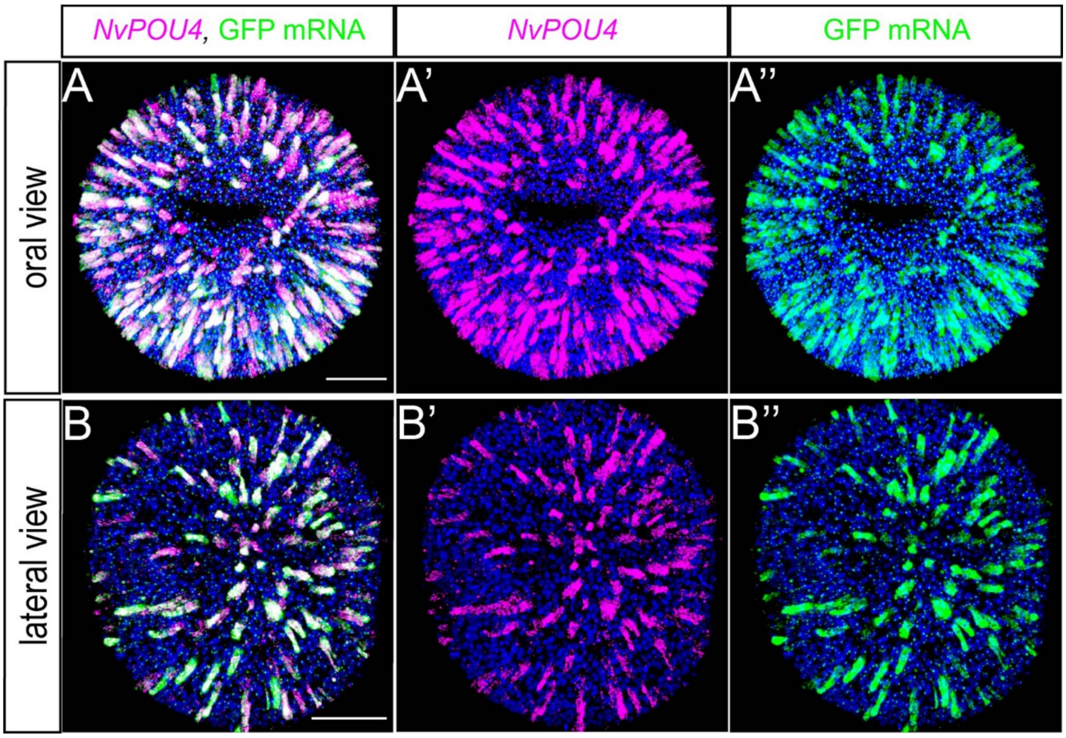


**Fig. S3 related to Fig. 3 Co-localization of *NvPOU4* and *memGFP* mRNA in transgenic embryos.** Double fluorescent *in situ*, using probes for *NvPOU4* (magenta), *memGFP* (green) and DAPI (blue), demonstrates that the reporter gene expression mimics endogenous *NvPOU4* expression at blastula stage in embryos from the *NvPOU4*::memGFP transgenic line. (A-A") oral views; (B-B") lateral views with the aboral pole to the left. Scale bars represent 50 µm

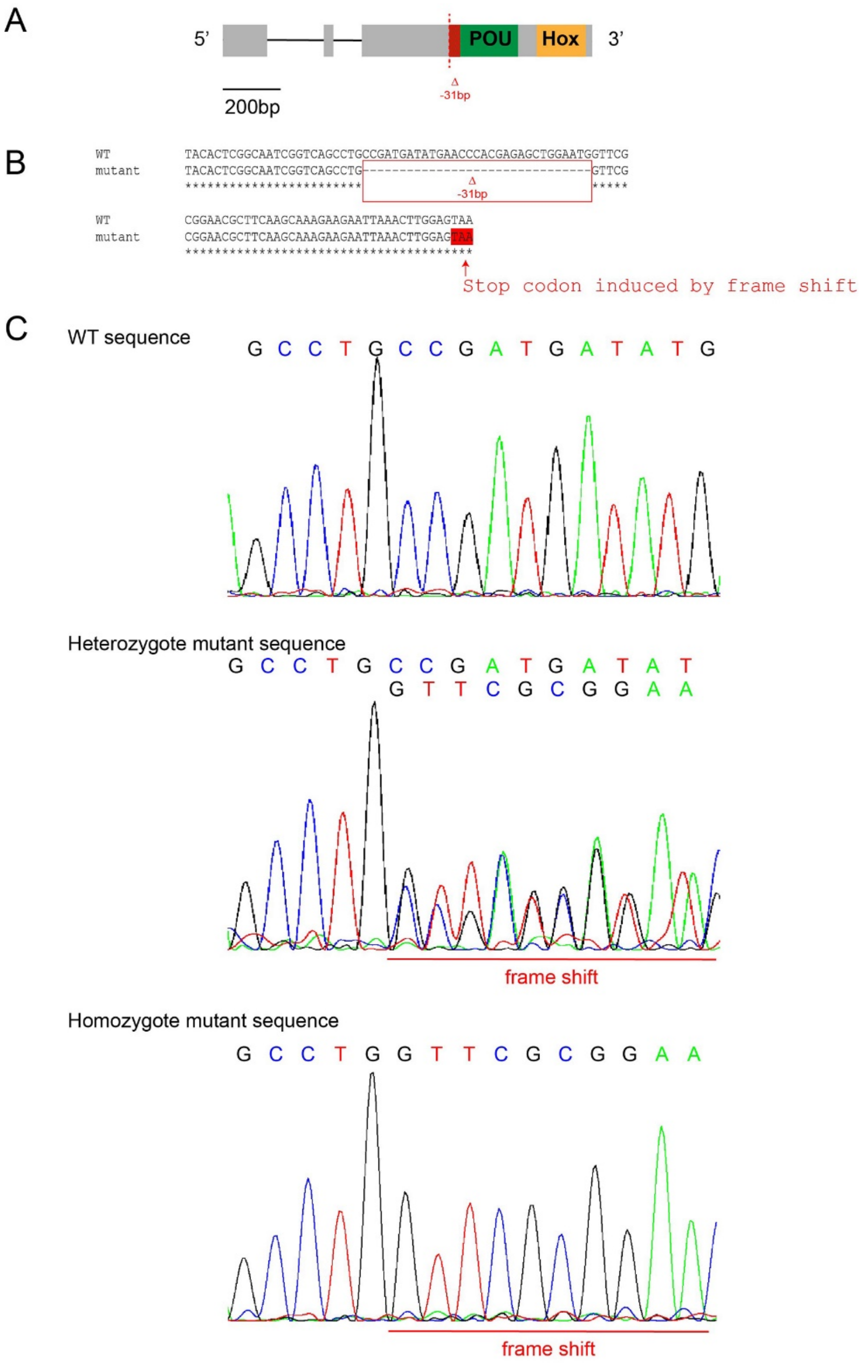


**Fig. S4 related to Fig. 5 Generation of *NvPOU4*$^{-/-}$ via CRISPR/Cas9 generate a 31bp deletion.** (A) Schematic of the knock out targeting strategy. Exons are in grey boxes, the POU domain is shown as a green and the homeodomain as a yellow box. The gRNA targets the start of the POU domain (red line) and generated a deletion of 31bp (red box) causing a frame shift and the appearance of a premature STOP codon. (B) Sequence alignment between the wild-type and the mutant sequences. The STOP codon is highlighted in red. (C) DNA chromatograms derived from individual animals with wildtype, heterozygous or homozygous *NvPOU4* mutant genotype.

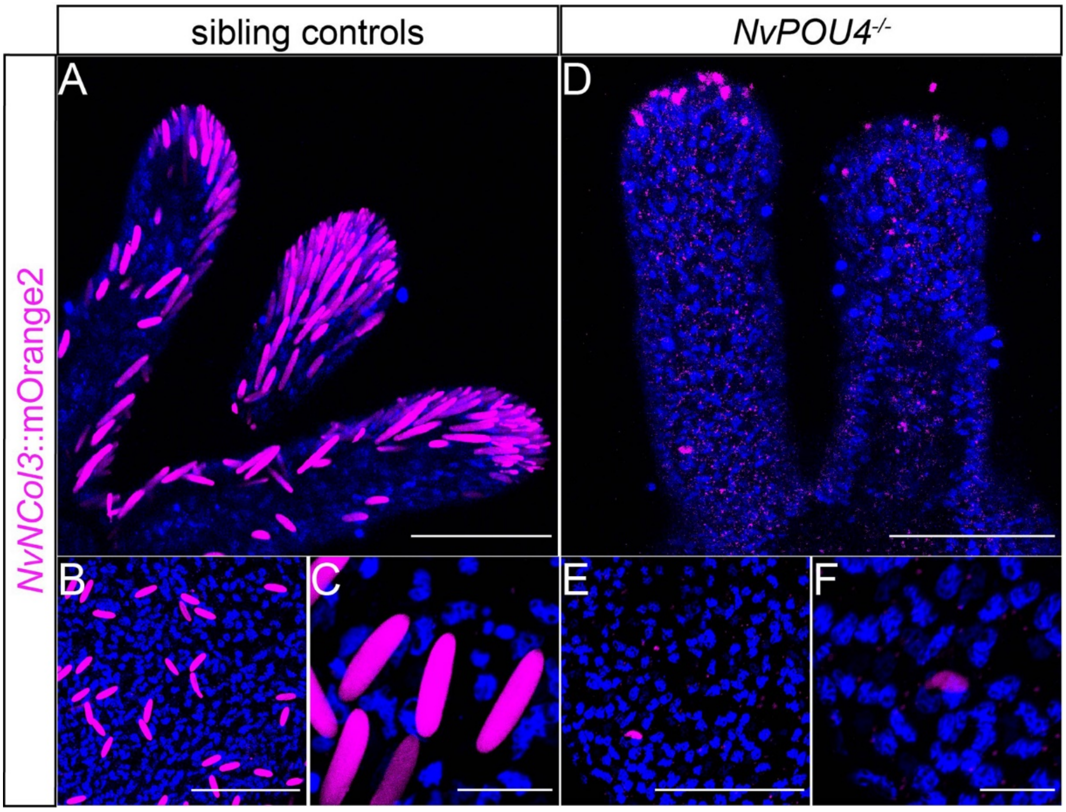


**Fig. S5 related to Fig. 5 Lack of mature cnidocytes in *NvNCol3*::mOrange2 transgenics mutant for *NvPOU4*.** (A-F) Antibody staining of mOrange2 protein (dsRed antibody, in magenta) and DAPI (blue) in (A-C) sibling controls (*NvPOU4*$^{+/+}$ or *NvPOU4*$^{+/-}$, *NvNcol3*::mOrange2$^{+/-}$) and (D-F) *NvPOU4*$^{-/-}$, *NvNcol3*::mOrange2$^{+/-}$ primary polyps. (A, D) tentacle tips, (B, E) body column, (C, F) higher magnification pictures of the cnidocyte capsules. *NvPOU4*$^{-/-}$ animals lack fully differentiated cnidocyte capsules, but still have some NvNCol3. Scale bar represents 50 µm (A-B, D-E) and 10 µm (C- and F), respectively.

A

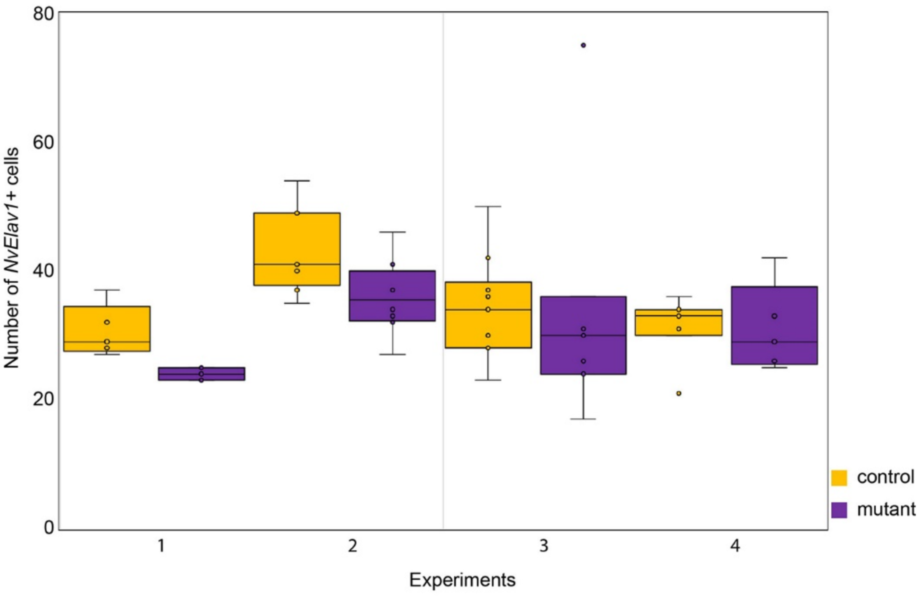


B

| | Experiment 1 | | Experiment 2 | | Experiment 3 | | Experiment 4 | |
|---|---|---|---|---|---|---|---|---|
| | control 1 | mutants 1 | control 2 | mutants 2 | control 3 | mutants 3 | control 4 | mutant 4 |
| | 32 | 23 | 54 | 33 | 42 | 36 | 30 | 29 |
| | 37 | 23 | 41 | 37 | 30 | 75 | 36 | 33 |
| | 29 | 24 | 35 | 46 | 34 | 17 | 21 | 42 |
| | 27 | 25 | 49 | 32 | 28 | 24 | 33 | 25 |
| | 28 | 25 | 37 | 41 | 37 | 26 | 34 | 26 |
| | | | 40 | 27 | 50 | 30 | 34 | |
| | | | 41 | 37 | 36 | 31 | 31 | |
| | | | 49 | 34 | 34 | | | |
| | | | | | 23 | | | |
| | | | | | 28 | | | |
| **median** | **29** | **24** | **41** | **35.5** | **34** | **30** | **33** | **29** |
| **average** | **30.3** | **24** | **43.3** | **35.9** | **34.2** | **34.1** | **31.29** | **31** |
| **standard dev** | **4.0** | **1** | **6.6** | **5.8** | **7.76** | **19.0** | **4.96** | **6.89** |
| **T Test** | | **0.01** | | **0.03** | | **0.98** | | **0.93** |
| **U test** | | | | | | **0.4** | | |

**Fig. S6 related to Fig. 6 The loss of *NvPOU4* in *Nematostella* does not affect the number of *NvElav1*$^{+}$ neurons** (A) Graphical representation showing the number of *NvElav1*$^{+}$ cells counted in a 100 µm x 100 µm square in *NvPOU4*$^{-/-}$ (purple) and in sibling controls (yellow) at primary polyp stage in four independent experiments. Numbers for the four experiments are shown in (B). A shapiro-wilk test was first calculated to ensure the normal distribution of each data. Then in case of normal distribution a T-test was performed (Experiments 1, 2, 4). In case of abnormal distribution a Mann Whitney test (U-test) was performed (Experiment 3). In all counting experiments there is a general tendency that *NvPOU4*$^{-/-}$ animals have less *NvElav1*$^{+}$ cells than their siblings controls (shown by the median), but the difference was not statistically significant for replicates 3 and 4. This suggests that the loss of *NvPOU4* in *Nematostella* has no or only a very mild effect on the specification of *NvElav1*$^{+}$ neurons.

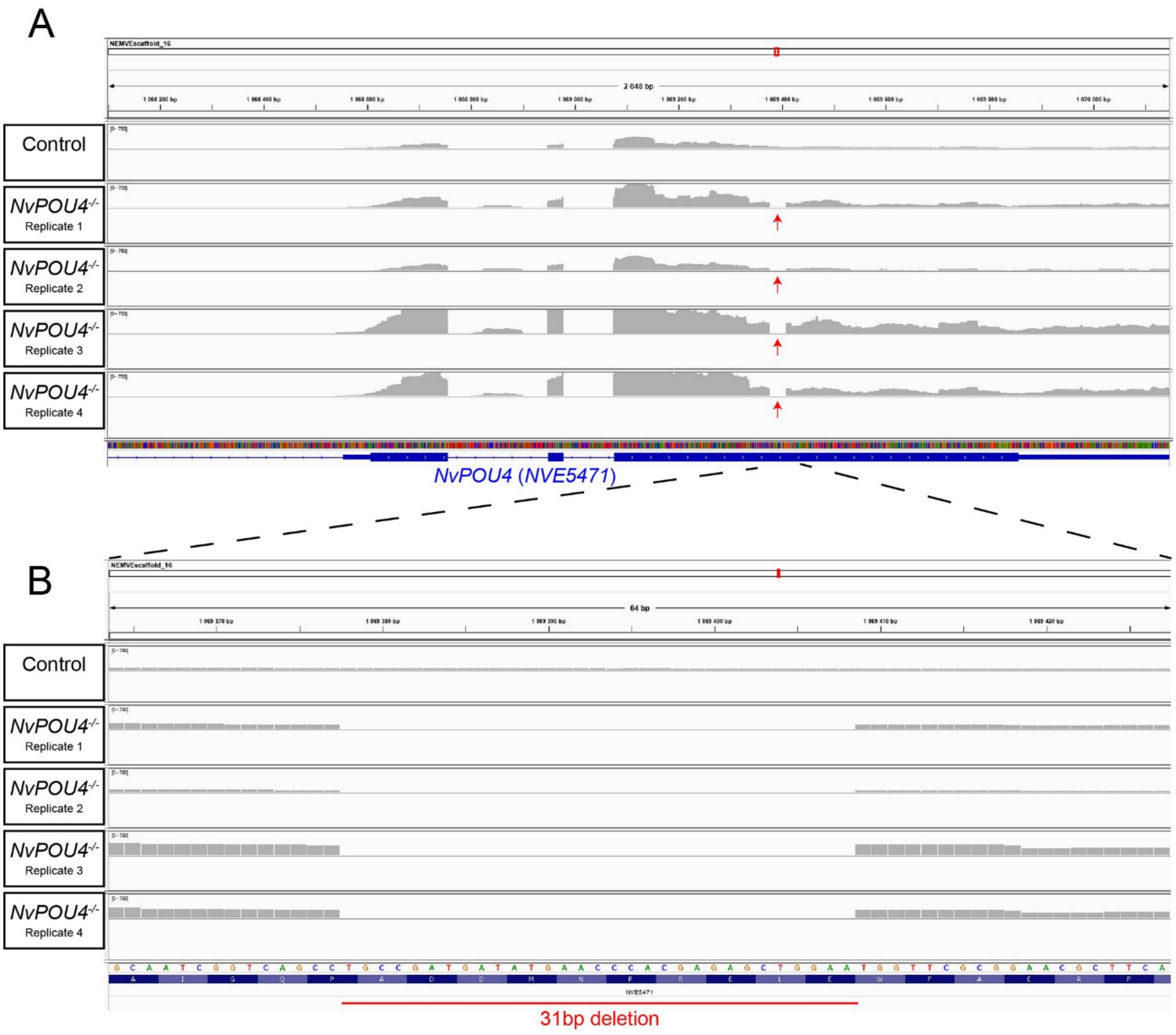


**Fig. S7 related to Fig. 6 RNA sequencing reads confirm the *NvPOU4* mutation at the mRNA level.** (A) RNA sequencing reads for the control and the four replicates of *NvPOU4*$^{-/-}$. In the mutants, no reads map to the deleted region (shown via a red arrow), whereas the control sample has reads mapping to it. (B) shows a higher magnification of the deleted region.

**Table S1** List of differentially expressed genes in *NvPOU4* mutants

**Table S2** Lists of genes differentially expressed in *NvPOU4* mutants and in *NvNCol3*::mOrange and *NvElav1*::mOrange transcriptomes

**Table S3** Transcription factors expressed in *NvPOU4*-positive metacells. Data derived from Table S4 in (Sebe-Pedros et al., 2018).

**Table S4** Overview of the crosses used in this study.

**Movie S1** Stack of double in situ hybridization for *NvPOU4* and *NvRFa* at blastula stage. *NvPOU4* in green, *NvRFa* in magenta, DAPI in grey. Related to Figure 2A.

**Movie S2** Stack of double in situ hybridization for *NvPOU4* and *NvRFa* at gastrula stage. *NvPOU4* in green, *NvRFa* in magenta, DAPI in grey. Related to Figure S2B.

**Movie S1** Stack of double in situ hybridization for *NvPOU4* and *NvRFa* at planula stage. *NvPOU4* in green, *NvRFa* in magenta, DAPI in grey. Related to Figure S2C.

**Movie S4** Stack of double in situ hybridization for *NvPOU4* and *NvNCol3* at blastula stage. *NvPOU4* in green, *NvNCol3* in magenta, DAPI in grey. Related to Figure 2C.

**Movie S5** Stack of double in situ hybridization for *NvPOU4* and *NvNCol3* at gastrula stage. *NvPOU4* in green, *NvNCol3* in magenta, DAPI in grey. Related to Figure S2E.

**Movie S6** Stack of double in situ hybridization for *NvPOU4* and *NvNCol3* at planula stage. *NvPOU4* in green, *NvNCol3* in magenta, DAPI in grey. Related to Figure S2F.

**Movie S7** Stack of double in situ hybridization for *NvPOU4* and *NvSoxB(2)* at blastula stage. *NvPOU4* in green, *NvSoxB(2)* in magenta, DAPI in grey. Related to Figure 2E.

**Movie S8** Stack of double in situ hybridization for *NvPOU4* and *NvSoxB(2)* at gastrula stage. *NvPOU4* in green, *NvSoxB(2)* in magenta, DAPI in grey. Related to Figure S2H.

**Movie S9** Stack of double in situ hybridization for *NvPOU4* and *NvSoxB(2)* at planula stage. *NvPOU4* in green, *NvSoxB(2)* in magenta, DAPI in grey. Related to Figure S2I.

**Movie S10** Stack of fluorescence in situ hybridization for *NvPOU4* and EdU incorporation at blastula stage. *NvPOU4* in green, EdU in magenta, DAPI in blue. Related to Figure 2G.
